# Development and characterisation of an optimised *in vitro* differentiation protocol for deriving hepatocyte-like cells from mouse embryonic stem cells

**DOI:** 10.64898/2026.05.13.724236

**Authors:** Beatrice Villani, Silviya Dimova-Vasileva, Abdulaziz Alhussini, Andrea Caporali, Congyue Chen, Ailsa Laird, C. Roland Wolf, Alistair Elfick, Richard R. Meehan, Sari Pennings

**Affiliations:** Institute for Neuroscience and Cardiovascular Research, College of Medicine and Veterinary Medicine, University of Edinburgh, Edinburgh, EH16 4TJ, UK; MRC Human Genetics Unit, Institute of Genetics and Cancer, College of Medicine and Veterinary Medicine, University of Edinburgh, Edinburgh, EH4 2XU, UK; Roslin Institute, University of Edinburgh, Easter Bush, Edinburgh, EH25 9RG; Institute for Regeneration and Repair Flow Cytometry and Cell Sorting Facility, College of Medicine and Veterinary Medicine, University of Edinburgh, Edinburgh, EH16 4TJ, UK; Department of Systems Medicine, School of Medicine, University of Dundee, Ninewells Hospital, Dundee, DD1 9SY, UK; Institute for Bioengineering, School of Engineering, University of Edinburgh, Kings Building, Edinburgh, EH9 3DW, UK

## Abstract

**Introduction:** Reliable generation of hepatocyte-like cells (HLCs) from pluripotent stem cells remains limited by heterogeneity and incomplete maturation of the cells. Derivation of induced pluripotent- and embryonic stem cells into hepatocytes typically relies on complex, and costly reagent-intensive protocols, with inconsistent reporting of differentiation efficiencies and functional maturation criteria. Variability in protocol designs highlights the need for optimisation, particularly in mouse embryonic stem cells (mESCs) systems that can be more comparable with mouse models for underpinning translational and toxicological studies. Here, we developed and evaluated two cytokine-based strategies: an advanced hepatic-inducing cocktail (A-HIC) and a simplified hepatic-inducing cocktail (HIC), both designed to reduce complexity while increasing functional maturation.

**Methods:** Hepatic differentiation and maturation were assessed by morphology, immunofluorescence, flow cytometry, and qRT-PCR. Functional competence was evaluated via urea production, glutathione synthesis, indocyanine green handling, cytochrome P450 inducibility, and impedance-based cell layer integrity monitoring.

**Results:** Morphological, molecular and phenotypic analyses confirmed that both protocols supported hepatic lineage progression, generating heterogeneous populations of hepatoblast-like and more mature HLCs. Gene expression confirmed the loss of pluripotency, transient endoderm induction, and subsequent hepatic specification. Functionally, cells exhibited glycogen storage, inducible urea production, glutathione depletion, and active ICG uptake and clearance, with stable monolayer formation by day 21. A-HIC-derived HLCs demonstrated enhanced maturation, with higher ASGR1 expression and stronger *Cyp1a1* induction.

**Discussion:** These findings suggest that both protocols generate functional HLCs; however, A-HIC yields a higher proportion of functionally mature cells with reduced variability. This approach enables a simple, cost-effective, and time-efficient generation of HLCs, supported by improved functional characterisation with potential applicability to more complex pluripotent systems, including human iPSC-based models for disease modelling and toxicology.

## Introduction

The liver has long been studied for its primary roles in drug metabolism, detoxification of xenobiotics and maintenance of systemic homeostasis, as well as its susceptibility to toxic injury (Mu et al., 2020; Liang et al., 2022). The use of mouse models has provided mechanistic insight into liver disease, hepatotoxicity of environmental and other toxins, and hepatic enzyme induction in response to pharmacological compounds (Sauer et al., 1997; Mu et al., 2020).

While anatomical differences exist between murine and human livers, most notably lobation in mice versus non-lobation in humans, both species share the same fundamental cell types, supporting cross-species comparison of core hepatic functions (Kruepunga et al., 2019). In both species, hepatocytes represent the dominant liver cell population, accounting for approximately 70% of liver cells, and are responsible for the majority of metabolic, detoxification, and homeostatic functions (Gao et al., 2008; Gong et al., 2022). Animal studies have advanced our understanding of liver development, physiology, and disease, but remain limited in their ability to accurately predict human drug responses (Barré-Sinoussi & Montagutelli, 2015). Ethics guidelines towards replacement, reduction, and refinement (3Rs) of animal use have reinforced the need for alternative, biologically relevant *in vitro* liver models with improved reproducibility and scalability for toxicological assessment and drug development.

Primary human hepatocytes (PHHs), considered the gold standard for toxicological assays, have disadvantages of limited availability, donor-dependent variability, high cost, and rapid loss of polarity and function in culture (Godoy et al., 2013; Korelova et al., 2019; Fu et al., 2023). Human hepatic cell lines such as HepG2 and HepaRG are widely used in toxicology and drug metabolism studies, however, they exhibit limited hepatic metabolic activity and low secretion of liver-associated proteins such as albumin (Ramaiahgari et al., 2014; Desai et al., 2017). Moreover, HepaRG cultures are costly to use under licence, while HepG2 carry extensive chromosomal alterations (Wiśniewski et al., 2016). Additionally, batch-to-batch variability and inconsistent inducer responses limit the predictive reliability of both cell lines (Henderson et al., 2019; Stanley & Wolf, 2022).

Pluripotent stem cells-derived hepatic cells have emerged as a promising alternative to primary hepatocytes for *in vitro* liver research, owing to their source of cells capable of self-renewal and differentiation into both parenchymal and non-parenchymal hepatic lineages (Fu et al., 2023). Induced pluripotent stem cells (iPSCs) have been shown to differentiate into hepatocytes and endothelial cells and to self-organise into three-dimensional spheroids and fibre-like structures (Miyamoto et al., 2015). These systems can generate multiple hepatic cell types, recapitulating liver cellular heterogeneity (Koui et al., 2017; Tasnim et al., 2019). Functionally, iPSC-derived hepatocytes have demonstrated improved sensitivity to hepatotoxic compounds compared with HepG2-based spheroids (Takayama et al., 2013). Some studies have shown toxicity profiles for tested compounds comparable to primary human hepatocytes but there is still debate regarding the reliability of iPSC-derived hepatic spheroids as predictive toxicity models (Lee et al., 2021; Yang et al., 2023).

Early efforts to generate hepatocyte-like cells from pluripotent stem cells were predominantly conducted using mouse embryonic stem cells (mESCs), reflecting their availability and genetic stability (Gadue et al., 2006; Kubo et al., 2004). These approaches attempted to recapitulate well-characterised murine developmental pathways *in vitro* but were limited by batch-to-batch variability in results (Smith et al., 1992; Conner, 2001; Lavon & Benvenisty, 2005). Several studies have established a range of stem cell differentiation protocols to generate hepatocyte-like cells from pluripotent sources (Hamazaki et al., 2001; Baharvand et al., 2008; Hay et al., 2007; Hay et al., 2008; Shiraki et al., 2008; Zhou et al., 2010; Cao et al., 2010; Q. Zhang et al., 2011; Pauwelyn et al., 2011; Ang et al., 2018). Across these studies, differentiation strategies typically followed a staged approach which, using the sequential addition of cytokines, recapitulates developmental signalling cues.

Firstly, cells are maintained under undifferentiated conditions by either the addition of Leukaemia Inhibitory Factor (LIF) or by culturing on mouse fibroblasts feeder layers (MEFs; Hamazaki et al., 2011; Baharvand et al., 2008; Zhou et al., 2010; Cao et al., 2010). Induction commonly involves Activin A-mediated endoderm specification, often combined with Wingless and Int-1 (WNT) or Bone Morphogenetic Protein (BMP) signalling modulation (Hay et al., 2007; Hay et al., 2008; Shiraki et al., 2008; Zhou et al., 2010; Pauwelyn et al., 2011; Ang et al., 2018). To drive cell commitment toward the definitive endoderm lineage, these protocols incorporated various fibroblast growth factors (FGF family members) but small changes in timing or dosage were seen to divert cells towards alternative lineages other than the liver (Haridoss et al., 2017; Shiraki et al., 2008; Engert et al., 2013; Touboul et al., 2010; Sumi et al., 2008). Similarly, later-stage maturation of the cells variably relies on addition of hepatocyte growth factor (HGF), sodium butyrate or dimethyl sulfoxide (DMSO) to promote hepatic lineage progression (Hamazaki et al., 2001; Zhou et al., 2010; Cao et al., 2010; Q. Zhang et al., 2011). Final maturation stages commonly relied on combinations of oncostatin M (OSM), dexamethasone (Dex), and insulin-transferrin-selenium (ITS) supplementation to enhance hepatocyte-associated phenotypes, further amplifying sensitivity to protocol variations (Li et al., 2010; Q. Zhang et al., 2011; R. Zhang et al., 2014; Chen et al., 2018; Demyanets et al., 2011; Magner et al., 2013; Shirahashi et al., 2004).

In all these protocols lineage commitment is governed through temporal and concentration-dependent signalling, but considerable variations exist in cytokine combinations, duration and media composition. Further protocol differences include the use of embryoid body formation versus adherent culture systems (Zhang et al., 2011; Shiraki et al., 2008) or feeder-dependent versus feeder-free conditions (Hamazaki et al., 2001; Hay et al., 2007). Some protocols restrict cells towards the hepatic fate by addition of inhibitory factors such as retinoid or BMP (Ang et al., 2018), whereas others use concentration changes of oxygen or foetal bovine serum (Pauwelyn et al., 2011). The increased use of human iPSC-derived hepatocytes has introduced new protocol variations using reprogramming factors (Reference).

All these studies demonstrated that mESCs, hESCs, and iPSCs, can be directed toward hepatocytic fates under defined conditions, although differentiation efficacy is still variable and reliant on complex and costly protocols (Kajiwara et al., 2012). Directly comparative data is lacking across systems, therefore there is a need to reassess and optimise protocols in mESC systems to provide a robust methodological framework for toxicological assessment across *in vivo* mouse models and *in vitro* hepatic differentiation models, which can inform approaches in human iPSC models and their translation to clinical studies.

In this study, we characterised and validated a simplified, cost-efficient and reproducible strategy for differentiating mouse embryonic stem cells (mESCs) into hepatocyte-like cells. We focused on reducing protocol complexity by minimising the number, concentration, and duration of cytokine and supplement exposure, while optimising the expression of key mature hepatic features relevant to toxicology and metabolic applications. Following extensive testing of protocol variations and further methodological development, the best two differentiation strategies, termed Hepatocyte-Inducing Cocktail (HIC) and Advanced Hepatocyte-Inducing Cocktail (A-HIC), were methodically evaluated during the differentiation process, with hepatocyte-like cell generation assessed using morphological, marker expression, and functional readouts (Figure 1). Our study advances new insights into the hepatocyte differentiation process and new methodology that will expand potential applications.

**Figure 1.**
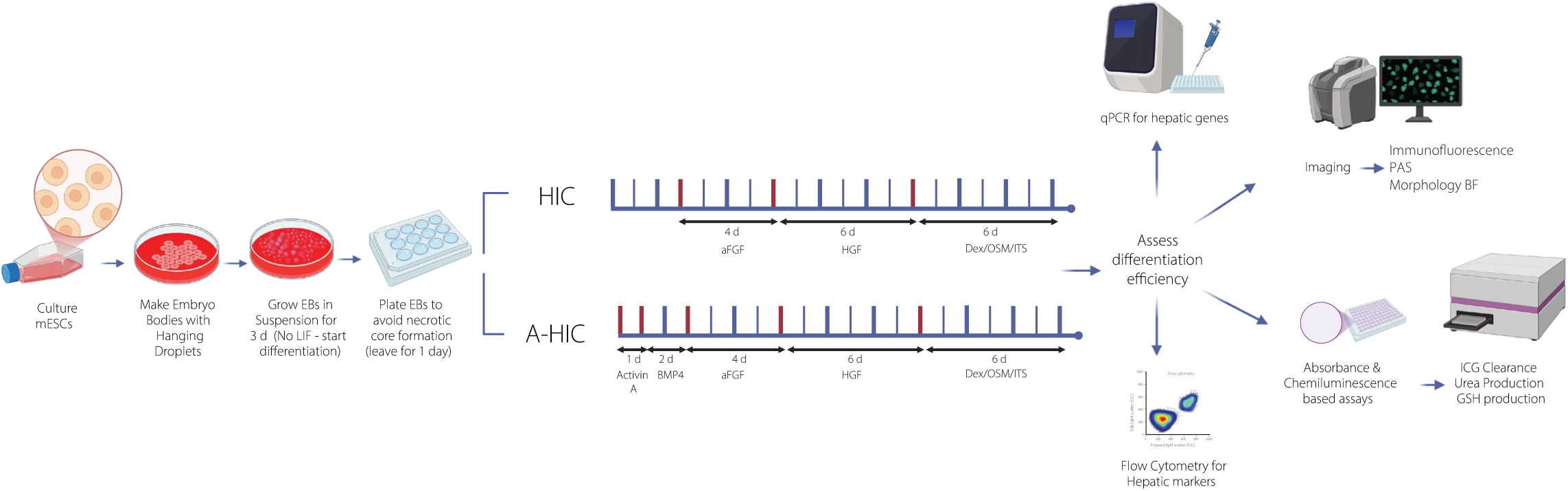
Graphical Abstract. Schematic overview of the A-HIC and HIC differentiation workflows, showing differentiation stages, cytokine exposure, and timing, together with the downstream morphological and functional readouts used for protocol evaluation. The workflows are presented on a ticked timeline, where each interval (|_|) corresponds to 1 day. Media was refreshed every 48 h. Bold ticks indicated media change events, with red bold ticks denoting transitions between cytokine conditions. Thin ticks represent periods during which cells were maintained without intervention.

## Methods

### Mouse pluripotent stem cells E14 culturing

ES-E14TG2a mouse embryonic stem cells (mESCs) were cultured on 0.2% gelatin-coated vessels prepared with porcine skin gelatin (Sigma-Aldrich, G1890). Cells were maintained under standard conditions (37 °C, 5% CO₂) in maintenance medium consisting of Dulbecco’s modified Eagle’s medium (DMEM with high glucose; Sigma, D5796) supplemented with 1% non-essential amino acids (Sigma, M7145), 1 mM sodium pyruvate (Sigma, S8636), 2 mM L-glutamine (Gibco, 25030-024), 12.5% foetal calf serum (Merck Millipore, F7524), 1x penicillin-streptomycin (Gibco, 15140122), 300 μM monothioglycerol (Sigma, M6145), 1000 u/mL recombinant mouse leukemia inhibitory factor (LIF; AMS Biotechnology, AMS-263-50), and 50 μM β-mercaptoethanol (Sigma, M3148). Cryopreserved cells were thawed and expanded for 2-3 passages under undifferentiated conditions prior to differentiation.

### Differentiation into hepatocyte-like-cells and cytokines reconstitution

All experiments for the study were done in triplicate unless otherwise stated. Cytokines and small molecules used for differentiation were reconstituted under sterile conditions according to manufacturer instructions. Activin A (20 ng/mL, #120-14E), Bone Morphogenetic Protein 4 (BMP4-10 ng/mL, #315-27), acidic Fibroblast Growth Factor (aFGF-50 ng/mL, #450-33A), Hepatocyte Growth Factor (HGF-20 ng/mL, #315-23), and Oncostatin M (OSM-10 ng/mL, #300-10) were prepared in sterile water or appropriate buffers, dexamethasone (Dex - 0.1 μM, #D8893) in ethanol, and insulin-transferrin-selenium (ITS) used at 1x. Cytokines were obtained from PeproTech, dexamethasone and ITS from Sigma-Aldrich. For differentiation, E14 mouse embryonic stem cells were dissociated using 0.25% trypsin-EDTA (Sigma, T4049) and used to generate embryoid bodies (EBs) by the hanging drop method (Zhou et al., 2005)(10 μL drops containing 10³ cells). EBs formed over 48 h and were subsequently cultured in suspension for an additional 3 days in medium lacking LIF). EBs were then plated onto 0.2% gelatin-coated culture wells and differentiated as outlined in Figure 1. More specifically, for protocol HIC, cells were maintained without added factors for the first 3 days, followed by treatment with aFGF (4 days, 50 ng/mL), HGF (6 days, 20 ng/mL) and a maturation phase of 3 days with OSM/Dex/ITS (10 ng/mL /0.1 μM/1X). For protocol A-HIC, cells were treated with Activin A (1 day, 20 ng/mL) followed by BMP4 (2 days, 10 ng/mL) prior to the same sequence of aFGF (4 days, 50 ng/mL), HGF (6 days, 20 ng/mL) and OSM/Dex/ITS (6 days, 10 ng/mL /0.1 μM/1X). For downstream analyses, EBs were seeded onto assay-specific substrates, including 12-well plates for RNA extraction, 24-well plates or μ-Slide chambers for imaging, and 96-well plates for metabolic assays. Impedance measurements were conducted using ECIS 8-well arrays (Applied Biophysics-Thistle Scientific, ABP-8W10E+PET).

### HepG2 cells culturing

HepG2 cells were used as comparator controls in differentiation and functional assays. HepG2 cells (kindly provided by Dr Shaden Melhem) were cultured at 37 °C, 5% CO₂ in DMEM with low glucose (Sigma, DD5546) supplemented with 15% foetal bovine serum (Sigma, F7524), 1% non-essential amino acids (Sigma, M7145), 1% L-glutamine (Gibco, 25030-024), 1% sodium pyruvate (Sigma, S8636), and 1% penicillin-streptomycin. Cells were passaged using 0.005% trypsin-EDTA (Gibco, 25200). For experimental assays, HepG2 cells were seeded at 8 x 10^4^-1 x 10^5^ cells per well in 12-well plates and 8 x 10^3^-2 x 10^4^ cells per well in 96-well plates.

### Immunocytochemistry

Immunofluorescence staining was performed directly in culture vessels (8-well μ-Slide chambers) or on sterile coverslips placed inside wells of 24 well plates. Embryoid bodies were fixed in 4% paraformaldehyde (15 min), washed in PBST (Tween 0.05%), and permeabilised with 0.4% Triton X-100 in PBS. Samples were blocked with 2% bovine serum albumin (Sigma, A2153) in PBST (1 h) and incubated with primary antibody at 4 °C overnight. This was followed by secondary antibodies (1:1000 in PBST, 1.5 h, dark) with 2 PBST washes between steps. Primary and secondary antibodies and working concentrations are listed in Table 1. Nuclei were stained with DAPI (1 μg/mL, Sigma, #P36934). Coverslips were mounted using ProLong Gold antifade mountant (Thermo Fisher Scientific, P36930). Images were acquired using a Nikon Ti-E inverted widefield fluorescence microscope and analysed with Volocity software.

**Table 1.** Antibodies used for immunocytochemistry and flow cytometric analyses.

| Antibody target | Host species | Manufacturer | Catalogue | Conjugation | Application | Working concentration / dilution |
| --- | --- | --- | --- | --- | --- | --- |
| HNF4a | Goat | Abcam | ab41898 | Unconjugated | Immunocytochemistry (IF) | 1:200 |
| Albumin (ALB) | Goat | Fortis Life Sciences | A90-134A | Unconjugated | Immunocytochemistry (IF) | 1:200 |
| DLK1 | Rabbit | Bioss | BS-2423R | Unconjugated | Immunocytochemistry (IF); Flow cytometry | 1:200 (IF) & 1:200 (FC) |
| AFP | Mouse | Bio-Techne | MAB1368 | Unconjugated | Immunocytochemistry (IF) | 1:200 |
| ASGR1 | Rabbit | Proteintech | 11739-1-AP | Unconjugated | Immunocytochemistry (IF); Flow cytometry | 0.40 μg per 1 x 10<sup>6</sup> cells |
| SSEA1 / CD15 | Mouse | BioLegend | 125610 | Alexa Fluor 488-conjugated | Flow cytometry | 5 μl per million cells in 100 μl staining volume |
| Donkey anti-goat IgG | Donkey | Thermo Fisher Scientific | A11057 | Alexa Fluor 568 | IF (secondary) Flow cytometry (secondary) | 1:1000 for IF/FC |
| Donkey anti-mouse IgG | Donkey | Thermo Fisher Scientific | A11057* | Alexa Fluor 488 | IF (secondary) & Flow cytometry (secondary) | 1:1000 for IF/FC |
| Donkey anti-rabbit IgG | Donkey | Thermo Fisher Scientific | A11057 | Alexa Fluor 568 | Flow cytometry (secondary) | 1:1000 for IF/FC |
| LIVE/DEAD™ Fixable Near-IR (780) | - | Thermo Fisher Scientific | L34994 | Near-IR dye | Flow cytometry (viability) | 1 μL per 1 x 10<sup>6</sup> cells |
| LIVE/DEAD™ Viability/Cytotoxicity Kit | - | Thermo Fisher Scientific | L3224 | Calcein AM / EthD-1 | Co-culture viability imaging | Manufacturer's instructions |

### RNA extraction, DNase clean-up and Reverse Transcription

Total RNA was isolated using RNeasy Mini Kit (Qiagen, 74104) with on-column DNase digestion (RNase-Free DNase Set, Qiagen, 79254) according to the manufacturer’s instructions. RNA was eluted in RNase-free water, quantified using a NanoDrop™ One spectrophotometer (Thermo Fisher Scientific), and stored at −80 °C. Complementary DNA (cDNA) was synthesised from 1 µg total RNA using the LunaScript RT SuperMix Kit (New England Biolabs, E3010S) following the manufacturer’s protocol (25 °C 2 min, 55 °C 10 min, 95 °C 1 min).

### Gene expression analysis with quantitative Polymerase Chain Reaction

Primer pairs were identified using validated sequences from the PrimerBank database (https://pga.mgh.harvard.edu/primerbank), synthesised by Eurofins Genomics (MWG, Germany), and resuspended in DNase/RNase-free water to 100 pmol/µL stocks. Primer performance was validated using standard curves generated from serial dilutions of tissue-specific cDNA under the qPCR conditions described above (Supplementary Figure S1). Reactions were performed using 100 nM forward and reverse primers. Primer sequences, amplification efficiencies, and correlation coefficients (R²) derived from the standard curves are listed in Table 2.

**Table 2.**
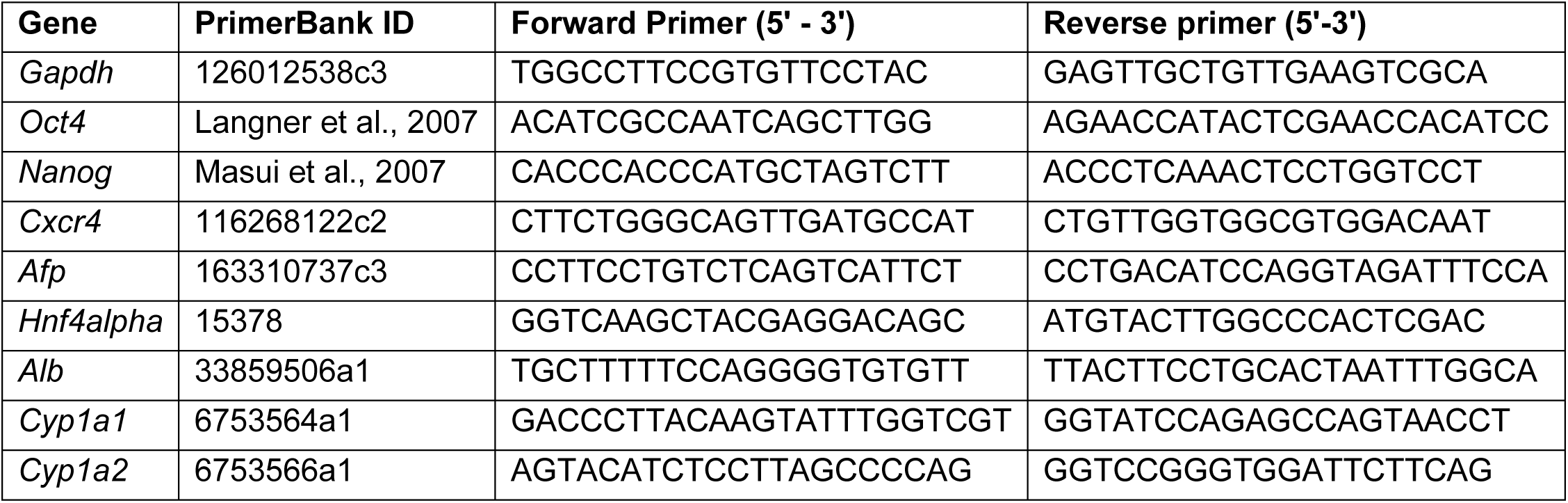
qPCr primers oligonucleotide sequences.

Quantitative real-time PCR (qPCR) was performed using PowerUp SYBR Green PCR Master Mix (Applied Biosystems, 4367659) in 10 μL reactions containing gene-specific primers (100 nM) and 1 μL cDNA template (equivalent to 25 ng RNA input). Reactions were assembled in MicroAmp™ Optical 384-well plates (Applied Biosystems, I309G0) and run on a QuantStudio 5 Real-Time PCR System. Thermal cycling consisted of 50 °C for 2 min, 95°C for 10 min, followed by 40 cycles of 95 °C for 15 s and 60 °C for 60 s. Melt curve analysis (95 °C - 60 °C - 95 °C) was performed to confirm amplification specificity. Gene expression of the gene of interest (GOI) was quantified using the ΔΔCt method, normalised to GAPDH (housekeeping gene), and expressed as fold change relative to time-matched control samples (day 0 or day 21) included on each plate using the following equation:

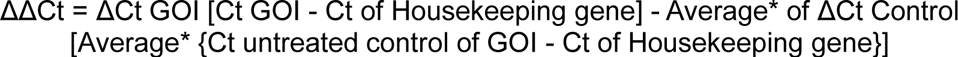

ΔΔCt-derived results are reported as fold change relative to the appropriate control, with associated 95% confidence intervals. Negative fold-change values indicate downregulation of transcript levels, whereas positive values indicate upregulation.

### Hepatic Nitrogen Cycle with urea production assay

At the completion of differentiation, culture supernatants were collected from hepatocyte-like cells (HLCs), as well as from HepG2 control cultures. Basal urea secretion was quantified using the QuantiChrom™ Urea Assay Kit (BioAssay Systems) according to the manufacturer’s instructions. To assess functional nitrogen metabolism, cells were additionally challenged for 24 hours with either 1 mM ammonium bicarbonate or 1 mM sodium bicarbonate, the latter serving as a control for possible increased alkalosis. Urea levels in supernatants collected following treatment were measured using the same assay. Absorbance readings were acquired at 520 nm using a VantaStar BMG plate reader.

### Flow cytometric profiling of hepatic lineage and maturation markers

HLCs cultured in 12-well plates were dissociated with trypsin-EDTA, pelleted (300 × g, 4 min), and resuspended in PBS. Cell viability was assessed using LIVE/DEAD™ Fixable Near-IR (780) dye (Thermo Fisher Scientific, L34994; 633 nm excitation) for 30 min at room temperature in the dark. Cells were washed, fixed in 4% paraformaldehyde (30 min), and washed in DPBS containing 0.1% Tween-20 (DPBS-T). Samples were permeabilised in 0.3% Triton X-100 (BioLegend, 39487) for 10 min, followed by primary antibody incubation overnight at 4 °C in the dark. After washing, cells were incubated with secondary antibodies (1:1000 in PBST) for 2 h on ice. Primary and secondary antibodies are listed in Table 1. Prior to acquisition, samples were washed, filtered to remove aggregates, and analysed using a NovoCyte Penteon flow cytometer (Agilent, 5-laser configuration). Negative controls included unstained cells, Live/Dead-only, secondary-only, and primary-only controls.

### Hepatic redox capacity with total glutathione (GSH/GSSG) quantification

HepG2 cells were used to optimise assay conditions and determine a working concentration of L-buthionine sulfoximine (BSO; Sigma, B2515), a competitive inhibitor of glutathione synthesis. A dilution series (100-12 μM) was prepared from a 50 mg/mL DPBS stock. Cells were harvested by brief trypsinisation, pelleted (1200 × g, 4 min), resuspended in maintenance medium, and seeded at 8,000 cells per well in 96-well plates (Corning, 3841). After 24 h incubation with BSO, intracellular glutathione (GSH) levels depletion was quantified using the GSH-Glo™ Glutathione Assay (Promega, V6911) following the manufacturer’s instructions. The optimised BSO concentration was then applied to HLCs differentiated via A-HIC and HIC using identical seeding densities and incubation conditions. Luminescence was measured using a VantaStar BMG plate reader (0.25 s integration).

### Indocyanine green (ICG) uptake and release assay

A 5 mg/mL stock of indocyanine green (ICG; USP, 1340009-100MG) was prepared in DMSO. HLCs, HepG2 cells (control), and undifferentiated mESCs (negative control) were incubated with 0.5 mg/mL ICG in cytokine-free high-glucose DMEM. Supernatants were collected at 5-180 min to quantify ICG release and analysed at 820 nm using a VantaStar BMG plate reader (0.25 s integration) against a standard curve. To assess intracellular accumulation independently of release, cells were incubated with ICG for 1 h, briefly blocked with foetal bovine serum (adapted from Tapparo et al., 2017), lysed in RIPA buffer (Sigma, R0278-50ML), and absorbance measured at 820 nm. Values were normalised to viable cell counts.

### Glycogen Storage with Period Acid Schiff’s Staining

Glycogen accumulation in HLCs generated using A-HIC and HIC, as well as in HepG2 cells, was assessed by periodic acid-Schiff (PAS) staining. Cells were seeded on 13 mm diameter glass coverslips in 12-well plates and processed using a PAS staining kit (Sigma, 395B) following a protocol adapted from Hui et al. (2017). Cells were washed with DPBS, fixed in 4% paraformaldehyde (10 min), incubated with 0.5% periodic acid (5 min) and Schiff’s reagent (15 min), and counterstained with haematoxylin (2 min). Adult C57BL/6 mouse liver sections (28 weeks old, kindly provided by Angus Comerford) served as positive controls.

Coverslips were mounted using ProLong™ Gold antifade reagent (Thermo Fisher Scientific, P36930), imaged using a Nikon inverted widefield microscope with Volocity software.

### Electrical cell substrate impeding sensing

Electrical cell-substrate impeding sensing (ECIS) was performed using an ECIS Theta System (Applied BioPhysics) with 8-well arrays (Applied Biophysics-Thistle Scientific, ABP-8W10E+PET). Arrays were prepared according to manufacturer’s instructions: electrodes were treated with 10 mM sterile L-cysteine for 15 min, washed twice with sterile water, coated with 0.2% gelatin, and further equilibrated (cell-free) with 200 μl of culture medium. One embryoid body (EB) was plated per chamber and differentiated directly on the arrays. Impendence and resistance were recorded at 4 kHz throughout differentiation to monitor cell attachment and monolayer formation.

### Statistical Analysis

Statistical analyses were performed using GraphPad Prism (version 10). Comparisons between the two differentiation protocols were conducted using unpaired Student’s t-tests. For experiments involving multiple conditions (treatment versus untreated across both protocols ± -ve/+ve controls), statistical significance was assessed using one-way analysis of variance (ANOVA) followed by Sidak’s multiple comparisons test. Data are presented as mean ± standard deviation (SD) or 95% Confidence Intervals, and differences were considered statistically significant at p < 0.05.

## Results

Comparative research of liver responses in mouse models and *in vitro* hepatocyte models requires up-to-date and optimised protocols for *in vitro* mouse hepatocyte differentiation. Our preliminary comparative evaluation used mouse embryonic stem cells to derive and experimentally test a comprehensive selection of 13 protocols. These included published methods for hepatocyte differentiation from either mouse or human pluripotent cells and adaptations and modifications based on literature research focusing on cytokine-based embryoid body-initiated methods versus adhesion culture in feeder-free conditions, as well as newer methodological variations using transcription factor cell programming or metabolic control (Zhou et al., 2009; Cao et al.,2010; Pauwelyn et al., 2011; Rothová et al., 2018; Supplementary Table 1). After applying selection criteria for cell viability, hepatic marker gene expression as well as reagent burden and cost (Supplementary Figure S2), the two best performing protocols, which we named HIC and A-HIC, were selected and further evaluated through a series of rigorous comparative tests.

### Cobblestone-like hepatocyte morphology is present by day 21 in A-HIC and HIC cultures

Differentiating embryoid bodies (EBs) were longitudinally imaged to monitor morphological changes during differentiation under the HIC or A-HIC protocol (Figure 2). Following adhesion to the culture substrate, EBs progressively lost their spherical architecture as peripheral cells attached and initiated aggregate flattening (black circles). During intermediate stages, cells at the basal surface became overlaid by cells on the opposite side, forming a transient dome-like structure, producing superimposition of opposite EB regions (red circle). With continued differentiation this dome collapsed, frequently generating acellular cavities and regions of cellular debris indicative of localised cell loss (black arrowhead). Necrotic subpopulations were commonly observed in thicker EB regions, consistent with limited nutrient and oxygen diffusion (orange circle). No contractile activity or morphologies consistent with cardiomyocytes or other mesodermal derivatives were detected under either condition. Concomitantly, cell morphology evolved from elongated spindle-shaped cells arranged in loose fibroblast-like networks (black arrow) to mixed populations containing polygonal epithelial-like cells (purple circle). These progressively organised into dense sheet-like structures displaying a characteristic honeycomb or cobblestone pattern. By late differentiation stages (day 21-28), both protocols reproducibly generated discrete clusters of tightly packed cobblestone-like cells (black squares).

**Figure 2.**
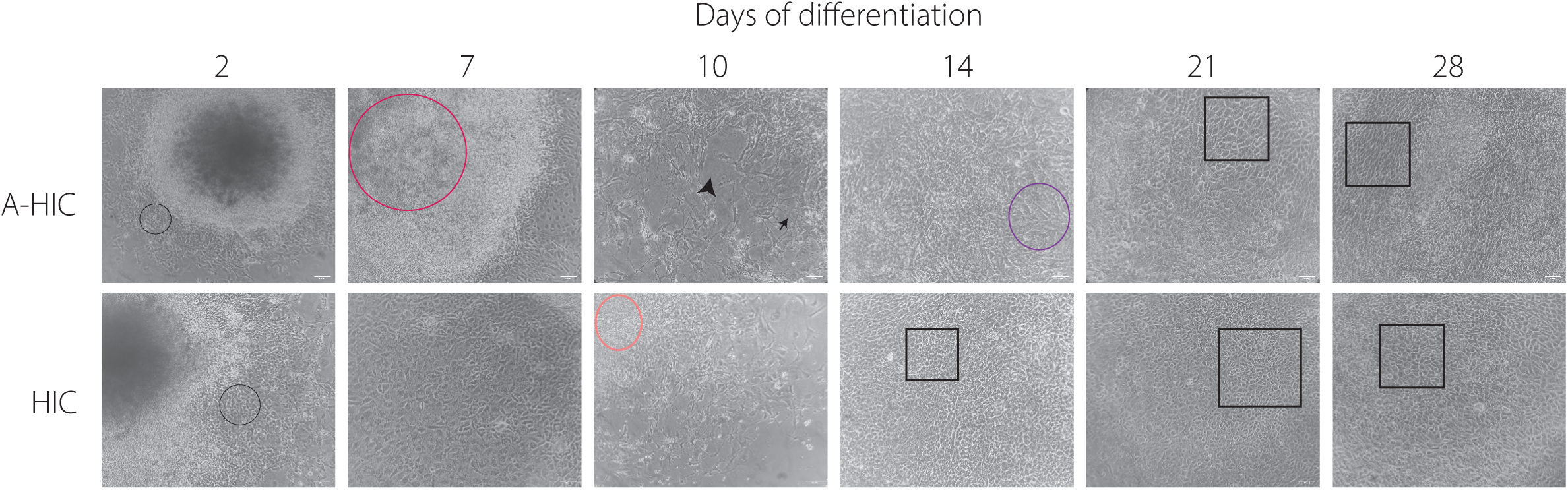
Representative brightfield images of hepatocyte-like cells differentiation using protocols A-HIC and HIC over 28 days. Figure 2 shows progressive morphological changes of distinct subpopulations within each protocol. Between day 2 and day 7, cells in contact with the surface, diffuse outward from the centre of the EB core. Black circles indicate progressive migration outwards from the EB core and loss of spherical architecture. The red circle indicates overlay formation of cells in the transient dome-like structure and superimposition of opposing EB regions. Between day 7 to 14, peripheral cells adopt a fibroblast-like (black arrow) morphology, while cells within the central regions retain polygonal epithelial-like cells (purple circle). Orange circle indicates the presence of necrotic subpopulations in thicker EB regions. Black arrowhead represents collapsed dome and presence of acellular cavity alongside loss of cell density. At later stages of differentiation (from day 14, but present in both protocols from day 21), both protocols generate populations of rounded, cobblestone-like cells that organise into compact, braid-like clusters, typical morphology adopted by hepatocyte cells in culture. These remain stable until day 28. Black squares correspond to cobblestone-like cells organised into clusters. Scale bars correspond to 100 µm.

### AFP, HNF4alpha and ALB expression indicates correct hepatic differentiation while revealing the presence of heterogenous subpopulations of HLCs

We first assessed hepatic lineage specification using detection of HNF4alpha, AFP, and ALB by immunofluorescence (Figure 3A-F). Qualitative analysis was undertaken due to the high cellular density within flattened EBs, resulting in pronounced multilayering and stratification, which restrict antibody penetration for optical sectioning and quantitative assessment. Accordingly, images are shown at both high and low magnification to capture subcellular localisation and overall signal distribution pattern and prevalence. Co-staining for HNF4alpha and AFP (Figure 3A) revealed discrete regions of partial overlap, with HNF4alpha being localised in the nuclei and AFP in the cytoplasm. As HNF4alpha expression is initiated during hepatic lineage commitment and maintained throughout hepatocyte maturation, the observed co-staining was consistent with hepatoblast and foetal-like hepatocyte populations.

**Figure 3.**
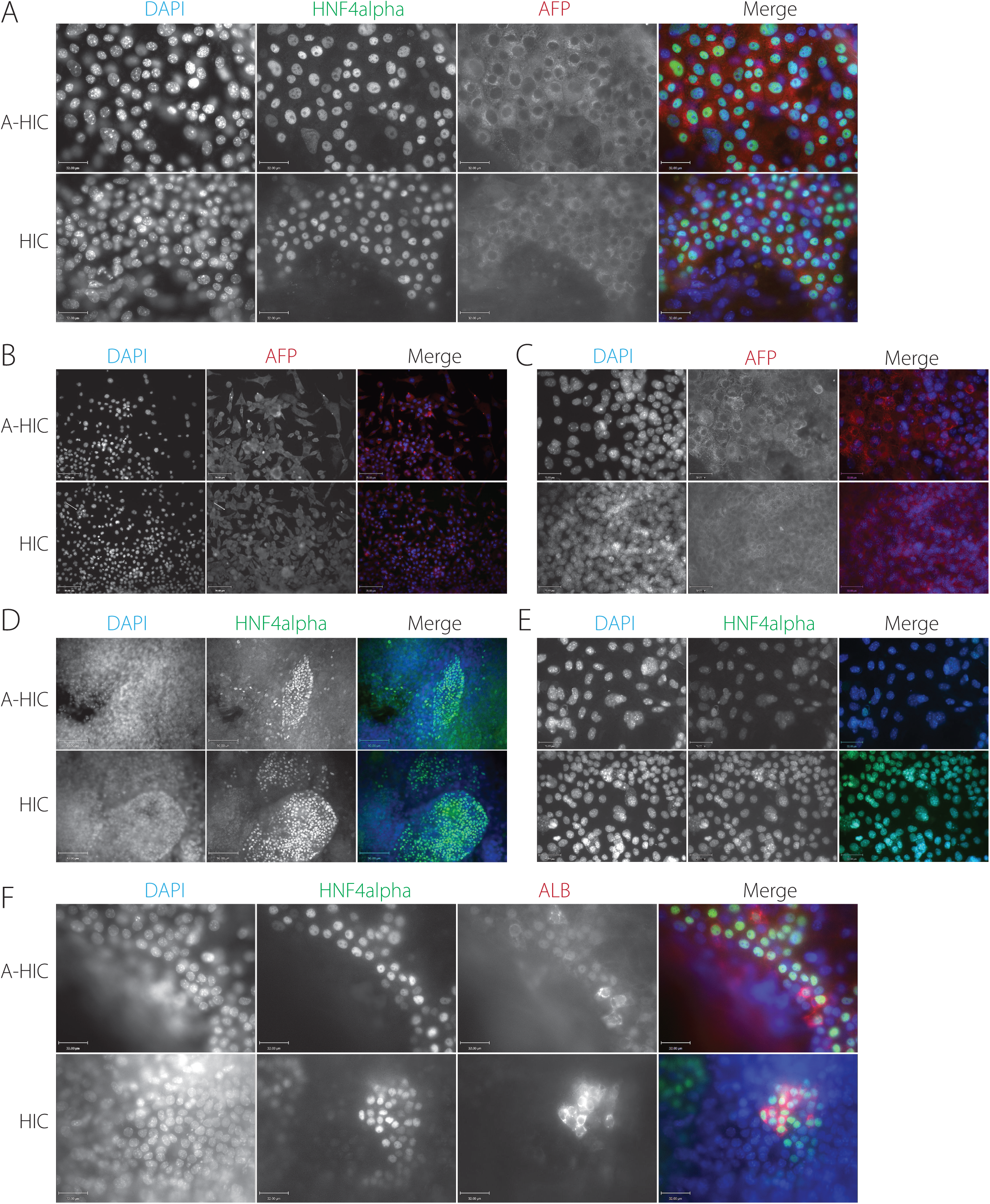
Characterisation of hepatic differentiation markers in hepatocyte-like cells (HLCs) generated using protocols A-HIC and HIC by immunofluorescence. Representative images show expression of the hepato-specific markers for both protocols at day 21 (end). **A)** Co-staining for HNF4alpha and AFP identifying transitional hepatoblast-foetal hepatocyte-like subpopulations. **B-C)** AFP expression (foetal hepatocyte marker) shown at (B) low magnification (scale bar: 70 µm), highlighting subcellular localisation, and (C) higher magnification (scale bar: 32 µm), illustrating spatial distribution and prevalence across the culture. White arrow, AFP negative cells. **D-E)** HNF4alpha expression (late hepatoblast/mature hepatic transcription factor) shown at (D) low magnification (scale bar: 90 µm), demonstrating nuclear localisation, and (E) higher magnification (scale bar: 32 µm), illustrating overall pattern and prevalence. **F)** Co-staining for HNF4a and ALB indicating progression toward a more mature hepatocyte-like phenotype. Images in panel (F) were acquired *in situ* through the culture plastic with cells maintained in PBS, whereas panels (B-E) were acquired following fixation and mounting of coverslips onto glass slides. Imaging clarity was affected by the plastic substrate (in situ) or dense cellular stratification inherent to the protocols (all images). Markers are shown in red or green, as indicated, with nuclear counterstaining by DAPI (blue). Scale bars: 32, 70 and 90µm.

Notably, AFP^+^ expression was consistently detected within HNF4alpha^+^ regions with no clear population of AFP^+^/HNF4alpha^-^ cells, suggesting that AFP expression occurred within HNF4alpha-expressing regions. To better resolve AFP signal and cellular distribution, single stainings for AFP (Figure 3B-C) and HN4alpha (Figure 3D-E) were performed. HNF4alpha-positive nuclei were predominantly observed in densely packed cell patches. In contrast, AFP-positive cells were detected in heterogeneous subpopulations, localised in the cytoplasm of elongated and polygonal cells arranged in networks or compact clusters.

Co-staining for HNFalpha (Figure 3F), expressed from early hepatocyte commitment to mature hepatocyte identity, and ALB, specific to mature hepatocyte identity, demonstrated that ALB expression was reduced relative to AFP, and appeared more spatially dispersed, with signal restricted to the cytoplasm. ALB and HNF4alpha did not consistently co-localise; while ALB⁺ cells were invariably HNF4alpha⁺, a subset of HNF4alpha⁺ cells lacked detectable ALB expression. For HN4alpha/ALB co-staining, additional lower-magnification images are provided in Supplementary Figure S3 to further illustrate signal distribution over larger areas. Overall, immunofluorescence observations were consistent with the presence of cells at distinct stages of maturation, supporting the existence of a mixed late-hepatoblast/hepatocyte-like population at day 21.

### DLK1 expressions confirms heterogenous transitioning hepatic subpopulations in A-HIC and HIC

To specifically investigate the presence of less mature, foetal like hepatocytes, DLK1 staining was performed (Figure 4A; Supplementary Figure S3B). DLK1 is a Delta-like 1 homolog transmembrane protein involved in liver development. Immunofluorescence imaging shows that DLK1 exhibits a punctate, predominantly membranous localisation consistent with a cell surface distribution. In addition to immunofluorescence localisation, this surface marker of foetal-like hepatocytes enables robust quantitative assessment of individual immunolabelled cells by flow cytometry. SSEA1, a pluripotency maker, was used as a control for undifferentiated cells. Flow cytometry analysis (Figure 4B) of DLK1/SSEA1 quadrant plots indicates that the majority of A-HIC cells occupied the DLK1^+^/SSEA1^-^quadrant (Figure 4B), whereas HIC cultures displayed greater dispersion with a trend toward a higher proportion of double-negative events compared to A-HIC cultures, although this did not reach statistical significance (p=0.093) due to inter-replicate variance in differentiation outcomes (9.80% ± 12.70 for A-HIC and 27.2% ± 19.10 for HIC, mean ± SD, Figure 4C). The proportion of DLK1^+^SSEA1^-^ cells between protocols (A-HIC: 89.47% ± 12.48 vs HIC: 72.15% ± 18.92, mean ± SD) did not differ significantly (Figure 4C).

**Figure 4.**
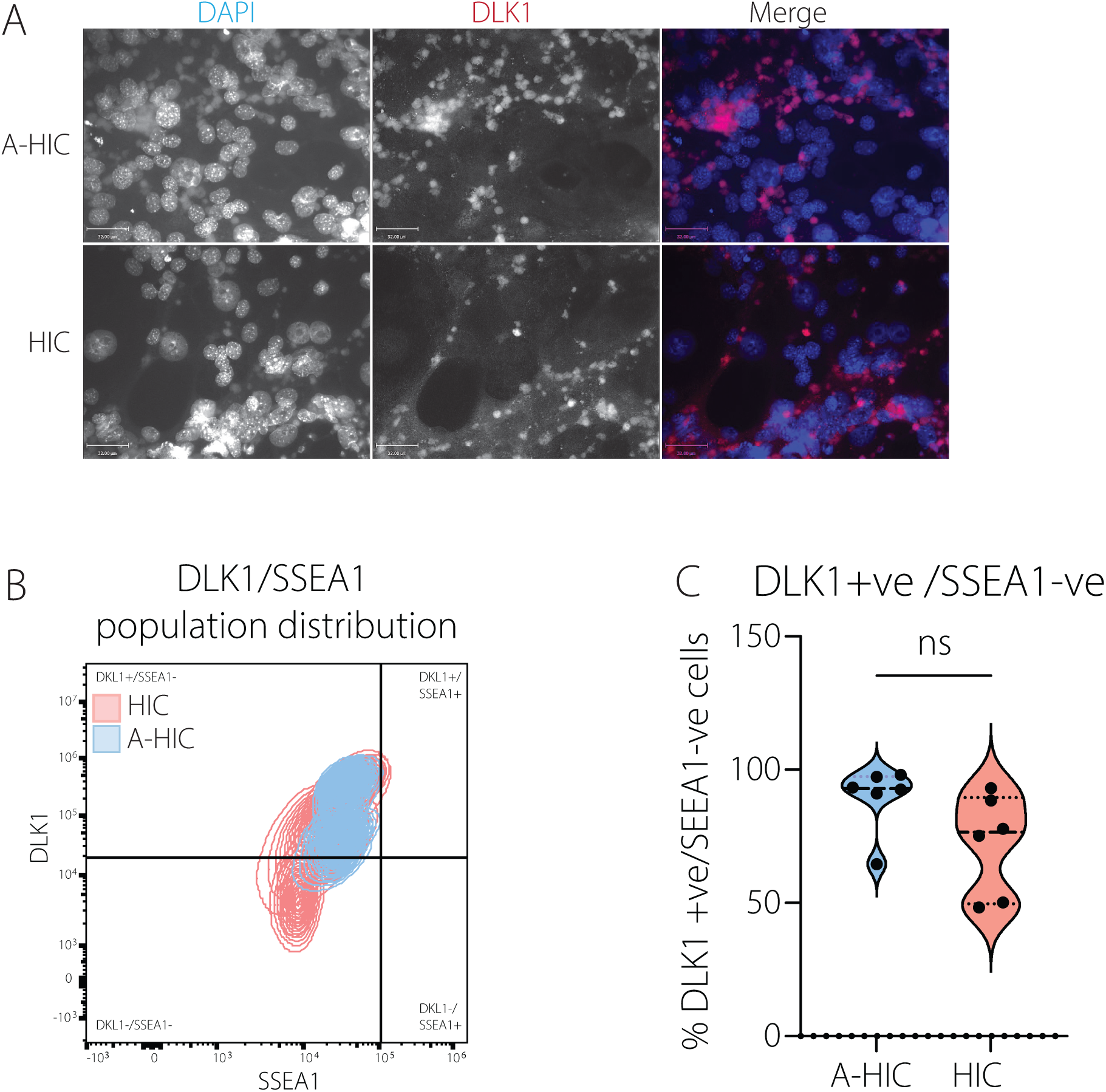
Assessment of Cellular heterogeneity in derived HLCs via DLK1 assessment. **A)** Representative images showing DLK1 expression (late hepatoblast-to-foetal hepatocyte marker). Marker is shown in red as indicated, with nuclear counterstaining by DAPI (blue). Scale bars: 32 µm**. B)** Representative DLK1 versus SSEA1 quadrant plots from juxtaposed samples. **C)** Quantification of DLK1+/SSEA1-cells across replicates for each protocol. Data are presented as individual replicates with violin distribution plots. Statistical analysis for panels was performed using Student t-test and it is indicated as *ns*, not significant.

### Transcript expression analysis confirms temporal progression from pluripotency to hepatic lineage in A-HIC and HIC cultures

To examine transcriptional changes during differentiation, quantitative PCR (qPCR) analysis was performed on cells differentiated using either protocol at days 0, 7, 14, 21, and 28 (Figure 5). Gene expression was normalised to day 0 and expressed as fold change (95% CI). From the differentiation trajectory outlined with stage-specific marker expression, a subset of representative genes was selected for targeted qPCR analysis (Figure 5A, genes in bold). Key stages of hepatic differentiation were assessed, including pluripotency markers (*Oct4, Nanog,* Figure 5B), meso-endodermal specification marker (*Cxcr4)* and early-to-late hepatoblast-to-foetal marker (*Afp)* (Figure 5C), early hepatocyte commitment to hepatocyte marker (*Hnf4alpha*) and hepatocyte maturation marker (*Alb*) (Figure 5D).

**Figure 5.**
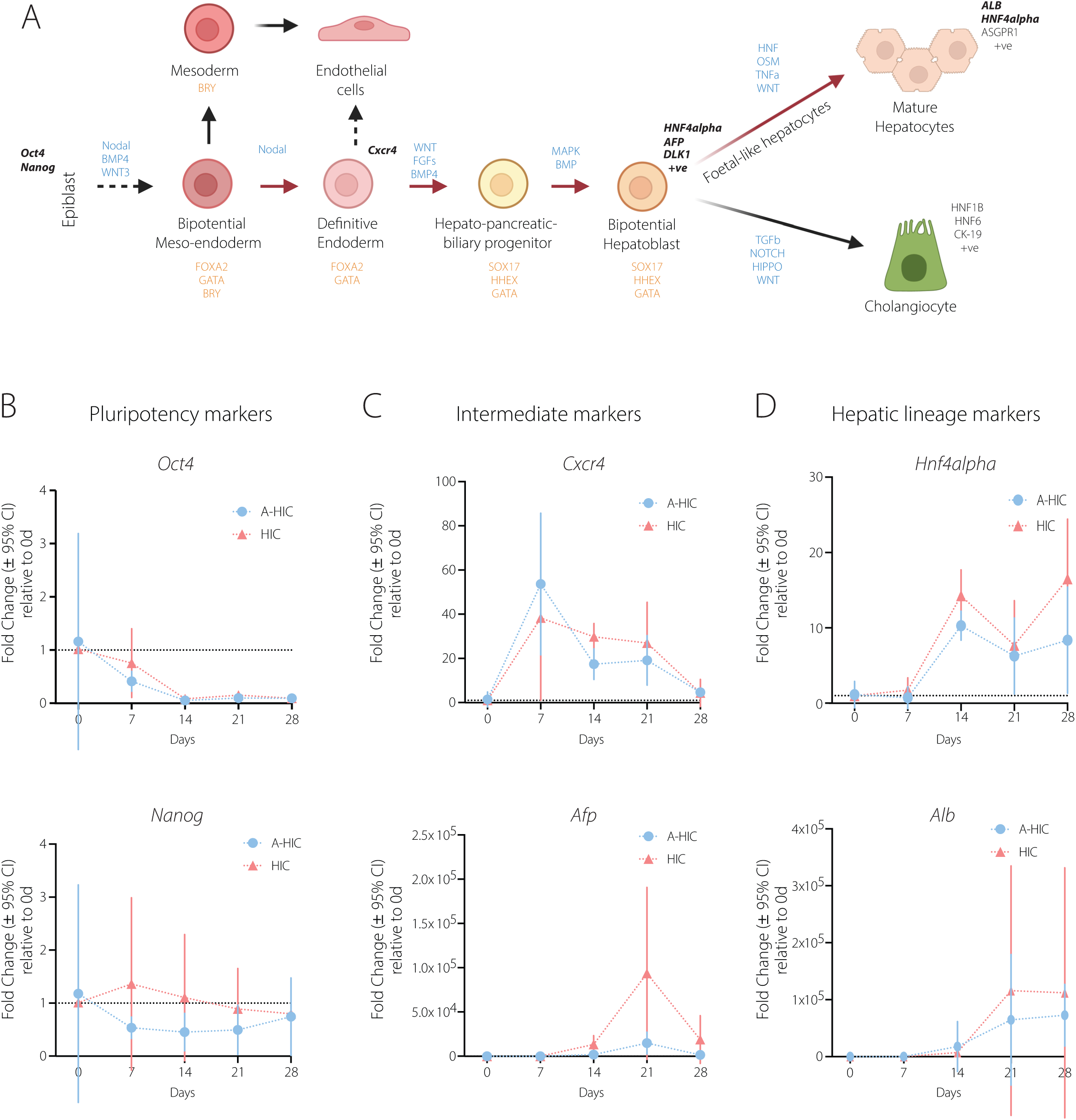
qPCR analysis of stage-specific gene expression during hepatocyte differentiation. **A)** Schematic representation of hepatic differentiation trajectories illustrating the intended progression followed by A-HIC and HIC protocols (red arrows), from epiblast through definitive endoderm and hepatoblast stages toward foetal-like and mature hepatocytes. Alternative lineage outcomes arising at intermediate stages are also indicated, including endothelial, mesenchymal, pancreatic, biliary, and cholangiocyte lineages. Active signalling pathways are shown in blue and stage-specific transcription factors in orange. Stage specific markers are in black, of those, selected qPCR gene markers are shown in bold. **B-D)** Gene expression was measured at days 0, 7, 14, 21, and 28 in hepatocyte-like cells generated using both differentiation protocols. **B**) Relative expression of pluripotency markers ***Oct4*** and ***Nanog*** during differentiation. **C)** Expression of intermediate meso-endodermal and hepatic progenitor markers ***Cxcr4*** and ***Afp.* D)** Expression of hepatic lineage markers ***Hnf4a*** and ***Alb***. Data is presented as fold change relative to day 0 (mean ± 95% CI, *n* = 3 biological replicates). The red dashed line indicates baseline expression (fold change = 1).

In both differentiation protocols, *Oct4* expression decreased markedly by day 14 and remained suppressed at later time points, indicating effective exit from pluripotency. *Nanog* expression followed a broadly similar trajectory. In A-HIC cultures, *Nanog* levels decreased markedly over time. In contrast, HIC cultures exhibited a modest upregulation up to day 7, followed by progressive downregulation from day 7 onwards. Across both protocols, *Nanog* expression was characterised by inter-replicate variability, consistent with heterogeneity within the differentiating populations*. Cxcr4* expression exhibited a substantial increase up to day 7 under both protocols, consistent with meso-endodermal specification. The magnitude of induction was variable between replicates. Thereafter, expression declined rapidly, in line with progression from definitive endoderm toward hepatocyte lineage specification and subsequent maturation, during which *Cxcr4* is downregulated. Genes associated with hepatic specification were detected at later stages. *Afp* expression remained low at early time points but increased at day 14, with a more pronounced induction observed under protocol HIC, suggesting the emergence of hepatoblast-like populations. For both protocols, *Afp* expression decreased past day 21. *Hnf4alpha* expression increased at intermediate stages in both conditions, specifically between day 7 and 14 and then again from day 21 to day 28, supporting activation throughout hepatic transcriptional programmes. In contrast, *Alb* expression was undetected up to day 14; thereafter it showed modest increase for both protocols until day 28. However, inter-replicate variability of these later genes was pronounced for both protocols, possibly indicating limited and heterogeneous progression towards mature hepatocyte identity. An analogous analysis was performed for *Cyp1a1* and *Cyp1a2* transcripts in unstimulated cultures across the same differentiation time points (days 0, 7, 14, 21, and 28) to characterise their baseline expression trajectories (Supplementary Figure S4). Together, these results suggest that both HIC and A-HIC differentiation protocols effectively drive exit from pluripotency and progression through meso-endodermal commitment toward hepatic lineage specification, although maturation remains variable and heterogeneous across replicates.

### Hepatic maturation is established by day 21 in A-HIC and HIC

To determine whether differentiation time could be reduced without compromising hepatic maturation or functional stability, we compared molecular and functional readouts at day 21 and day 28. qPCR analysis revealed minimal differences in hepatic gene expression between the two time points across the entire gene panel (Figure 5). Across both protocols, expression of pluripotency markers (*Oct4, Nanog*), the meso-endodermal marker (*Cxcr4*), hepatic lineage markers (*Afp, Hnf4alpha, Alb*) and main detoxifying genes (*Cyp1a1, Cyp1a2*) did not differ significantly between day 21 and day 28 (Supplementary Figure S5A-D). Basal urea production was also detectable at both stages with no significant differences (Supplementary Figure S5E). Together, these findings indicate that hepatic identity and basal metabolic function are largely established by day 21, supporting its selection as the optimal end differentiation time point.

### Hepato-specific functions are operational in A-HIC and HIC differentiated HLCs

Next, the functional maturity achieved by protocols A-HIC and HIC was assessed with a panel of complementary assays to evaluate key hepatic functions in live cells: glycogen storage, nitrogen metabolism, redox capacity, and substrate uptake and clearance.

### Glycogen storage via PAS staining in differentiated HLCs

Glycogen storage capacity was evaluated by periodic acid-Schiff (PAS) staining of HLCs generated using protocols A-HIC and HIC, compared with adult mouse liver and HepG2 cells as controls (Figure 6A). A-HIC-derived HLCs displayed strong magenta PAS staining with intensity and distribution comparable to adult mouse liver, indicating substantial intracellular glycogen accumulation. In contrast, HIC-derived HLCs showed weaker and more heterogeneous staining, suggesting reduced glycogen storage capacity. HepG2 cells exhibited little to no detectable PAS signal and formed multilayered epithelial aggregates with cells growing on top of one another and leaving discernible intercellular spaces. Overall, these findings suggested that protocol A-HIC provides HLCs with better glycogen storage.

**Figure 6.**
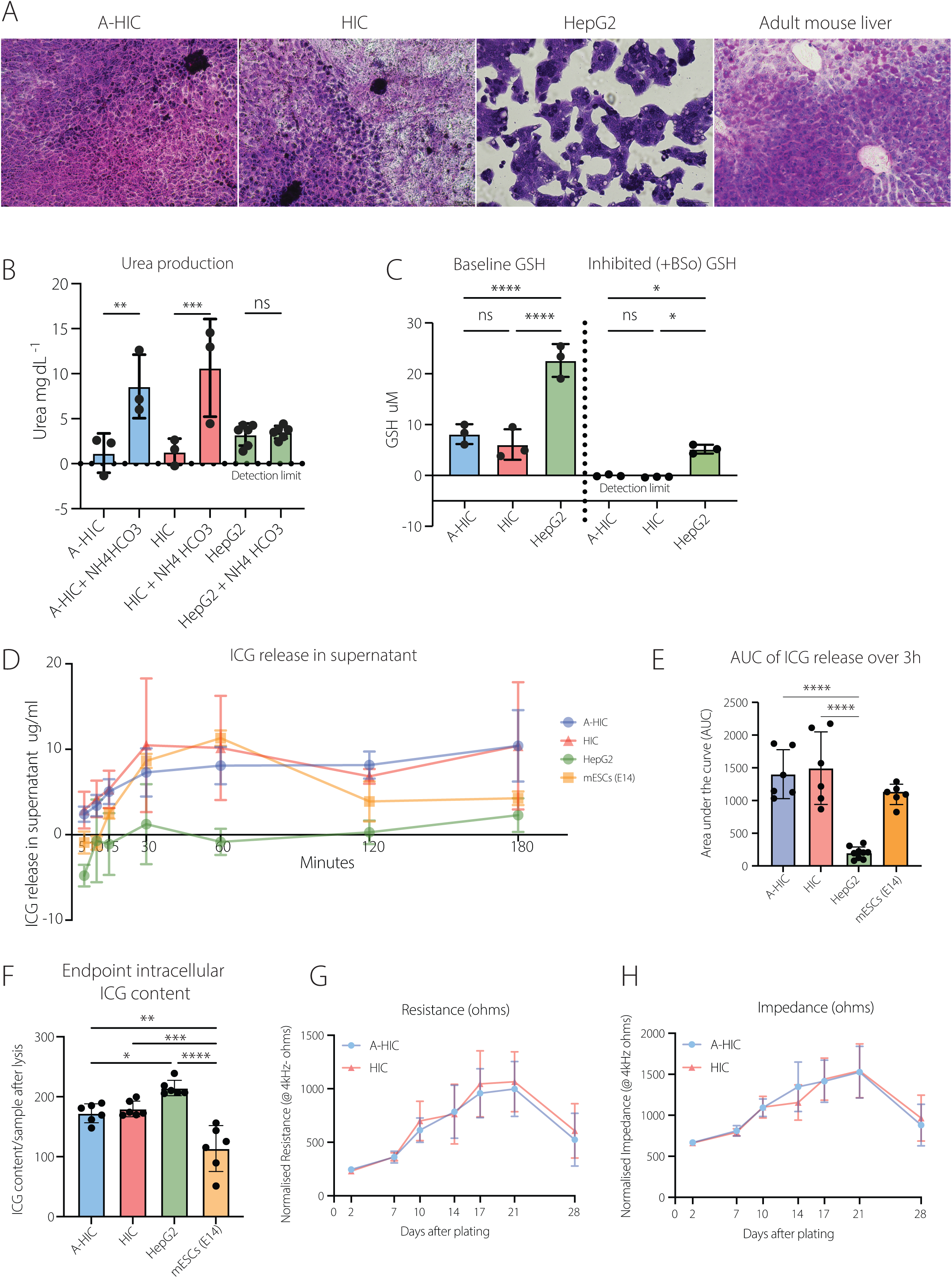
Comparative assessment of hepatic function in hepatocyte-like cells generated using A-HIC and HIC protocols. **A)** Representative images of glycogen storage assessed with Periodic acid-Schiff (PAS) staining for protocol A-HIC and HIC; HepG2 cells and adult mouse liver (28 weeks) were included as reference controls. **B)** Urea production measured under basal conditions and following stimulation with ammonium bicarbonate (NH₄HCO₃). **C)** Glutathione (GSH) depletion following exposure to buthionine sulfoximine (BSO). **D)** Hepatocyte-selective indocyanine green (ICG) release over time following ICG exposure, measured in A-HIC (blue), HIC (red) and compared with HepG2 cells (control, green) and undifferentiated mouse embryonic stem cells (-ve control, mESCs-E14, yellow). ICG concentrations (µg/mL) were calculated from absorbance measurements at 820 nm using linear regression (y = mx + c) derived from standard curves prepared for ICG in the corresponding culture media for each cell type. Only the linear range of each standard curve was used for quantification. **E)** Total ICG release quantified as the area under the curve (AUC) from time-resolved release profiles. **F)** Intracellular ICG retention quantified following cell lysis to assess endpoint dye accumulation. **G - H)** Electric cell-substrate impedance sensing (ECIS) analysis over a 28-day differentiation period showing representative resistance (G) and impedance (H) traces following plating onto ECIS electrodes. Data are presented as individual biological replicates with mean ± standard deviation (SD). Statistical analysis for panels (B, C, E, F) was performed using one-way ANOVA with Sidak’s multiple-comparisons post hoc test. Statistical significance is indicated as *ns*, not significant, *p ≤ 0.05, **p ≤ 0.01, ***p ≤ 0.001, ****p ≤ 0.0001.

### Urea production in differentiated HLCs

Next, hepatic nitrogen metabolism (Figure 6B) was assessed to determine whether differentiated cells could upregulate urea production in response to metabolic stimulation. Urea levels were measured under basal conditions and following 24-hour exposure to ammonium bicarbonate, a substrate expected to drive urea cycle activity in functional hepatocytes. In both HIC and A-HIC differentiation conditions, ammonium bicarbonate treatment resulted in a significant increase in urea production compared with untreated controls, indicating inducible urea cycle activity. Average urea levels were comparable but varied across biological replicates, reflecting differences in differentiation efficiency between experiments. HepG2 cells were analysed in parallel as a reference line and exhibited relatively stable basal urea production but showed no significant urea production increase following ammonium exposure, indicative of an altered nitrogen metabolism.

### Glutathione depletion following BSO exposure in differentiated HLCs

Cellular redox capacity was assessed by quantifying intracellular reduced glutathione (GSH) using the luminescence-based GSH-Glo assay (Figure 6C). GSH levels represent a balance of usage and synthesis to maintain normal antioxidant defence, whereas changes in intracellular GSH are widely used as indicators of oxidative stress, as GSH depletion can precede redox imbalance, impaired detoxification, and cell death (Liu et al., 2022). To determine whether differentiated cells could deplete GSH in the absence of GSH synthesis, cultures were treated with buthionine sulfoximine (BSO), an inhibitor of γ-glutamylcysteine synthetase. A dose-response analysis identified 25 µM BSO as an effective inhibitory concentration (Supplementary Figure S6). Under basal conditions, both A-HIC and HIC HLCs exhibited detectable GSH levels, which were significantly reduced following BSO treatment, indicative of active redox capacity maintained by glutathione synthesis. HepG2 cells displayed higher basal GSH levels and retained measurable GSH after BSO exposure. Collectively, the data indicates that HLCs generated by both protocols possess functional glutathione biosynthesis and redox responsiveness, consistent with an active and regulatable detoxification pathway.

### Total and Intracellular ICG uptake and release in differentiated HLCs

Hepatic uptake and clearance capacity were assessed using indocyanine green (ICG), quantified through time-dependent release in live cultures (Figure 6D-E) and intracellular accumulation following cell lysis (Figure 6F). ICG is a clinically used dye that is taken up by hepatocytes via organic anion transporting polypeptides (OATPs) and excreted unchanged through the canalicular transporter MRP2; therefore, its handling reflects hepatocellular transport activity. Two reference models were included for comparison. Undifferentiated mouse embryonic stem cells (mESCs) served as a negative control, while HepG2 cells were used as a hepatocyte reference due to their widespread use in functional assays. ICG concentrations (µg/mL) were calculated from absorbance measurements at 820 nm by spectrophotometry, using standard curves generated in the corresponding culture media for each condition (Supplementary Figure S7A-C), applying linear regression within the validated linear range. Hepatocyte-like cells generated using both differentiation protocols showed progressive ICG release into the medium over the 3-hour monitoring period, consistent with active uptake and efflux. Total ICG release was quantified as the area under the curve (AUC) of concentration-time profiles (μg/ml x min) derived from the data shown in Figure 5D. AUC values were comparable between protocols (A-HIC: 1402 ± 373; HIC: 1495 ± 553, mean ± SD), with no significant difference detected between HIC and A-HIC conditions. Undifferentiated mESCs displayed lower extracellular ICG levels (1094 ± 155) and a transient increase during early time-points followed by decline. However, viability analysis demonstrated a substantial reduction in cell viability, indicating that extracellular dye likely resulted from membrane disruption rather than functional transport (Supplementary Figure S7D). In contrast, time-point measurements and integrated total AUC quantification for HepG2 cells showed minimal ICG excretion (199 ± 90.6), indicating limited expression and/or functionality of MRP2 canalicular transporters required for ICG efflux. To assess ICG uptake independently of extracellular clearance, intracellular ICG accumulation was quantified following cell lysis using an approach adapted from Tapparo et al. (2024; Figure 6F). Both A-HIC and HIC-derived HLCs showed low intracellular ICG retention at endpoint, consistent with efficient export over time. In contrast, HepG2 cells exhibited significantly higher intracellular ICG levels compared with HLCs. mESCs used as a negative control confirmed minimal intracellular ICG accumulation in non-hepatic cells. Collectively, these findings suggest that HIC and A-HIC HLCs exhibit functional hepatocellular transport activity via uptake and clearance, while HepG2 cells indicate impaired efflux, and mESCs show non-specific dye uptake and release due to reduced viability.

### Longitudinal analysis of cell layer integrity via resistance and impedance in differentiating HLC cultures

Electric cell-substrate impedance sensing (ECIS) was used to monitor cell attachment, spreading, and monolayer stability over 28 days of differentiation. Resistance (Figure 6G) and impedance (Figure 6H) increased during the first 10-14 days following plating, reaching a plateau of approximately 1000-1500 Ω corresponding to stable electrode coverage and cohesive cell layers. These strong barrier properties are indicative of the formation of a functional epithelial layer with matured tight cell junctions (Meng & Roy, 2016). After approximately day 21, both values declined, coincident with observed overgrowth and consistent with reduced epithelial integrity. The data suggest that differentiating cultures under HIC or A-HIC conditions establish stable, confluent layers by day 10-14, with maximal integrity maintained up to approximately day 21, after which overgrowth compromises barrier function and cell-substrate interactions.

As ECIS analysis suggested progressive establishment of stable cellular monolayers over time, we additionally assessed whether culture on PET membrane substrates affected maintenance of hepatic identity, as substrate composition can influence cell attachment, polarity and maintenance of differentiated phenotypes (Kress et al., 2012).

Immunofluorescence analysis confirmed retention of hepatic identity following transfer to PET membranes, with discrete HNF4alpha-positive clusters and variable ASGR1 expression comparable to standard plate- and flask-based cultures (Supplementary Figure S8).

### A-HIC exhibits enhanced hepatocyte maturation and xenobiotic responsiveness compared to HIC

#### Functional hepatocyte maturation levels in differentiated HLCs

The previous characterisations suggested heterogeneous differentiating populations of cells (Figure 3). To assess the efficiency of hepatic maturation achieved by each differentiation protocol, HLC cellular composition was quantified by flow cytometry (Figure 7A-D) using the gating strategy shown in Supplementary Figure S9. Populations generated using A-HIC and HIC were analysed for ASGR1, a marker for mature hepatocyte-like cells and glycoprotein homeostasis functionality. ASGR1 is a major subunit of the asialoglycoprotein receptor mediating the endocytosis and lysosomal degradation of glycoproteins. ASGR1 fluorescence histograms showed a larger shift in A-HIC samples relative to HIC, consistent with enrichment of ASGR1-expressing cells (Figure 7A). A-HIC cultures contained a significantly higher fraction of ASGR1^+^ cells compared with HIC-derived populations (90.25 ± 17.49% vs 55.09 ± 34.09%, Figure 7B). Despite variability across independent differentiations, A-HIC consistently generated populations enriched in ASGR1-expressing cells, indicating a more reproducible progression toward a mature hepatocyte-like functional phenotype.

**Figure 7.**
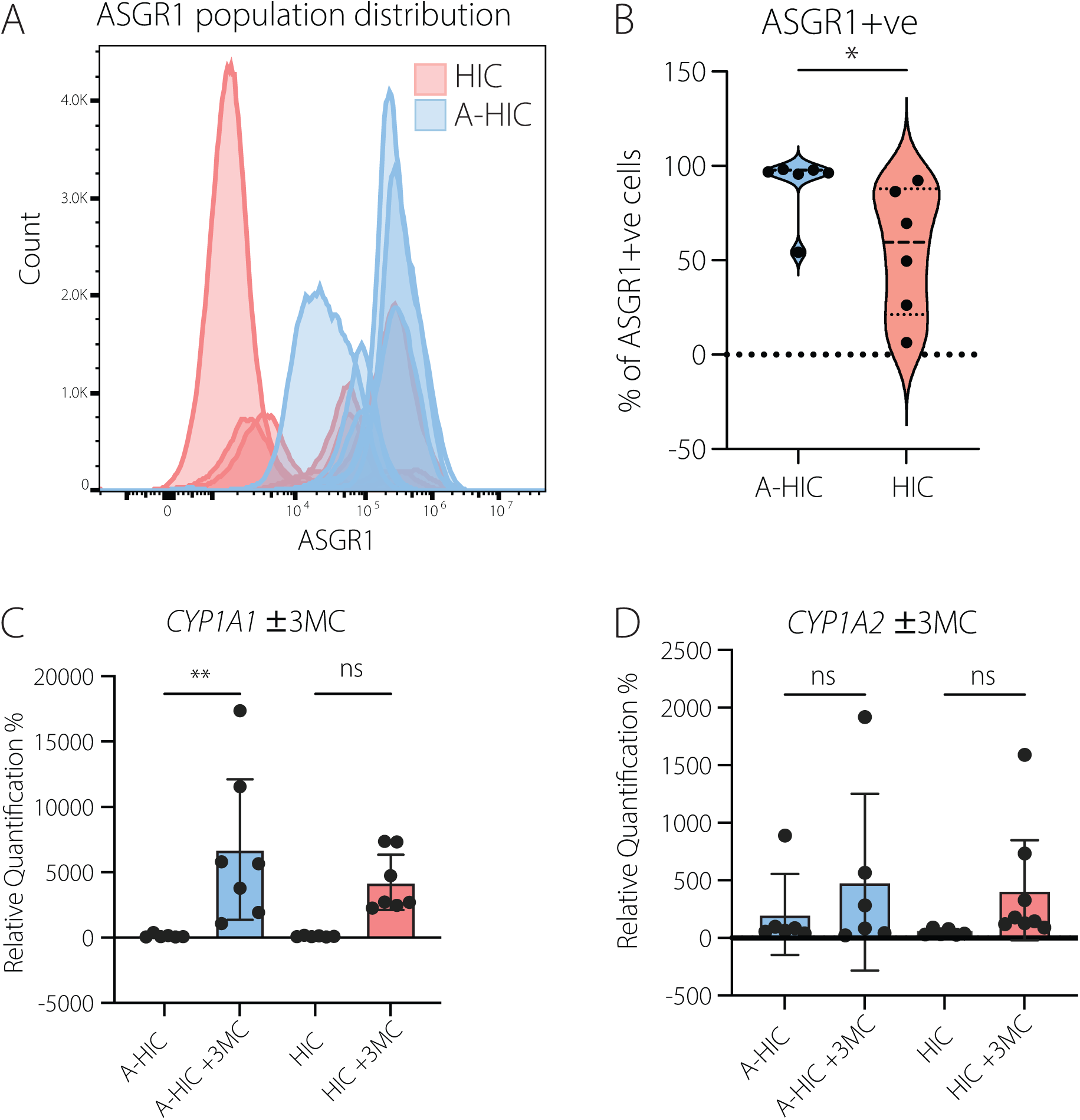
Quantification of mature hepatocytes-like cells and detox capacity. **A)** Representative overlaid histograms showing ASGR1 fluorescence distributions for individual replicates of A-HIC (blue) and HIC (red) cultures. **B)** Quantification of ASGR1^+^ cells across replicates for each protocol. **C-D)** *Cyp1a1 (*E) and *Cyp1a2* (F) mRNA levels quantified in both protocols by qPCR after 24 hrs exposure to 3-methylcholanthrene (3MC, 1 µM) and expressed relative to vehicle-treated controls. Data are presented as individual replicates with violin distribution plots; Statistical analysis for panels was performed using Student t-test and it is indicated as *p ≤ 0.05 (B). For C-D, data are presented as individual biological replicates with mean ± 95% confidence intervals. Statistical analysis was performed using one-way ANOVA with Sidak’s multiple-comparisons post hoc test and it is indicated as **p ≤ 0.01 or *ns*, not significant (C & D).

### Xenobiotic responsiveness in differentiated HLCs

To evaluate their inducible xenobiotic response, HLCs differentiated under each protocol were exposed to a prototypical cytochrome P450 inducer, 3-methylcholanthrene (3-MC; Jiang et al., 2009), and *Cyp1a1* and *Cyp1a2* mRNA expression was measured (Figure 7C-D). Following 24-hour exposure to a minimal dose (1 µM) of 3MC, A-HIC-derived HLCs exhibited upregulation of *Cyp1a1* transcripts (p=0.0043). The magnitude of induction was variable, with notable inter-replicate heterogeneity; however, an overall increase relative to vehicle controls was observed. In contrast, HIC-derived cells did not demonstrate a statistically significant increase in *Cyp1a1* expression compared to their respective vehicle-treated controls, despite a modest upward trend (p=0.0964). For *Cyp1A2*, neither differentiation protocol showed significant transcriptional induction following 3MC exposure relative to vehicle controls due to inter-replicate variability in the response (p=0.6925 for A-HIC & p=0.4395 for HIC).

Taken together, these findings suggest that the A-HIC protocol promotes a more consistent maturation toward a hepatocyte-like phenotype, as evidenced by enrichment of ASGR1⁺ cells and selective inducibility of CYP1A1, and suggested by other criteria we assessed, whereas HIC-derived populations remain more heterogeneous and exhibit limited xenobiotic responsiveness. Having established A-HIC as the best protocol for HLC differentiation, the evaluation assays also showed the functional robustness of these cells for downstream applications.

## Discussion

In this study, we derived two new differentiation protocols, named HIC and A-HIC, and evaluated their ability to direct murine embryonic stem cells toward a hepatocyte-like phenotype. We assessed their differentiation efficiency through cell morphological, molecular, and functional characterisations, ultimately revealing the best protocol.

Importantly, the differentiation methods described here were developed based on an initial comparative evaluation of 13 protocols, testing classic mouse differentiation methods as well as adapting to newer strategies including those used in human hepatocyte differentiation. We addressed the need for an updated mouse protocol fit for developing in vitro approaches that can be directly comparable to mouse model research. Our aim was to design a simplified and cost-effective protocol, minimising reliance on elaborate multi-step induction methods or complex media compositions. This consideration is particularly relevant given that many current differentiation protocols for pluripotent stem cells, including both ESCs and iPSCs, rely on combinations of growth factors, small molecules, and specialised media supplements, which substantially increase cost and limit scalability for applications (Deng et al., 2019).

Based on liver marker expression criteria, hepatocyte-inducing cytokine cocktails (HIC) emerged as more efficacious in preliminary testing, of which the two best performing variants were rigorously tested. Across the range of assays performed in this study, both differentiation protocols successfully directed mESCs toward a hepatocyte-like phenotype. Although the overall differentiation trajectory was broadly comparable between the two approaches, cultures generated using A-HIC displayed more consistent outcomes across independent experiments and multiple assays in achieving functionally mature hepatocyte characteristics.

For instance, flow cytometric analysis indicated that A-HIC cultures contained a higher proportion of cells progressing toward a mature hepatocyte-like phenotype. In contrast to many published mESC to HLCs differentiation protocols that reported predominantly on immunofluorescence across small fields of cultured cells, we additionally used multiparametric flow cytometry to quantify marker expression, providing a more representative estimate of the total yield of hepatocyte-like cells in each condition. Using DLK1/SSEA1 to define hepatic progenitors and ASGR1 as a mature hepatocyte surface marker, our A-HIC protocol reproducibly generated a highly homogeneous population, with most cells DLK1⁺SSEA1⁻ and more than 90% ASGR1⁺ with lower inter-experimental variability.

By comparison, in other studies it is often difficult to assess the true proportion of fully mature hepatocyte-like cells in the culture. An mESC protocol using sodium butyrate and cytokines reported efficient differentiation, but evaluation of maturation markers by flow cytometry could further support the role of the short-chain fatty acid in the context of its broader epigenetic effects on cell phenotype (Zhou et al., 2010). Another study reported that 71% of cells expressed coagulation factor VIII as a gene-therapy proof-of-concept but quantitation of canonical mature hepatocyte markers at the single-cell level could increase this method’s applicability (Cao et al., 2010). Many published protocols use immunofluorescence or semi-quantitative analyses of HNF4alpha or albumin expression to provide valuable spatial information on the differentiation state of the culture (Chen et al., 2010, Pauwelyn et al., 2011). The insight gained from this study is that these commonly used markers and methods were not sufficient to determine functional hepatocyte maturation, as our tested protocols’ benefits were only revealed in assays and markers for hepatic function, which were not available for most protocols in current usage.

A previously noted, many mESC-to-hepatocyte differentiation protocols follow a broadly embryonic developmental framework relying on complex culture systems. In one such approach, effective mouse hepatocyte maturation was promoted by co-culture with human liver non-parenchymal cell lines, exposure to hepatocyte growth factor derivatives and dexamethasone, and reporter-based sorting (Soto-Gutierrez et al., 2007). In contrast, the A-HIC protocol achieves hepatic specification within a simpler EB-based framework demonstrating hepatic commitment and maturation across the culture population and a more quantitatively robust route to generating larger, homogeneous populations of mature hepatocyte-like cells.

Importantly, cells differentiated using protocol A-HIC demonstrated the capacity and sensitivity to respond to xenobiotic stimulation following exposure to a low concentration of the aryl hydrocarbon receptor agonist 3-MC. Although induction varied between replicates, A-HIC cultures showed upregulation of *Cyp1a1,* indicating activation of hepatocyte-associated detoxification pathways, whereas HIC-derived cells did not show significant induction. It should be noted that the concentration of 3-MC used in this study was intentionally kept low to avoid cytotoxicity rather than to achieve maximal induction, which may partly explain the variability observed across replicates (Chowdhury et al., 2017).

Additional metabolic readouts were consistent with these findings. A-HIC cultures demonstrated more consistent glycogen accumulation within the cytoplasm and showed reduced inter-replicate variability compared with HIC-derived cultures. The cell surface expression of ASGR1 indicated more glycoprotein homeostasis functionality. Taken together, these observations suggest that the A-HIC conditions may promote more reproducible differentiation outcomes and support the emergence of a larger proportion of cells displaying mature hepatocyte characteristics.

One of the main limitations of these differentiation protocols was the degree of variability observed between independent replicates across both protocols. Inherent heterogeneity is commonly reported in differentiation systems and underscores the importance of replication and the use of complementary functional assays. Although chemically defined media have improved reproducibility, differentiation efficiency and functional maturation remain influenced by media composition, oxygen tension, nutrient availability, and cytokine timing and dosage (Hay et al., 2008; Shiraki et al., 2008; Si-Tayeb et al., 2010; Jung et al., 2016; Boon et al., 2020). In mitigation this study aimed to adopt a simplified differentiation protocol to minimise additional potential sources of experimental variability. Our preliminary comparative testing suggested that the EB-based approach is justified by better performance Despite more than two decades of research, the generation of functionally mature hepatocyte-like cells from pluripotent stem cells remains challenging and highly sensitive to protocol design and experimental context (Graffmann et al., 2022). The HIC protocol used in this study represents a modified adaptation of the differentiation protocol originally described by Hamazaki et al. (2001). This protocol was selected as it performed well as one of the earlier and relatively simple approaches for directing mESC differentiation toward hepatocyte-like cells, providing a useful baseline against which more complex strategies could be evaluated. Modifications were introduced to improve practicality and reproducibility. Culture conditions were simplified by reducing the number of media formulations, relying primarily on high-glucose Dulbecco’s Medium and not including the original protocol’s dual use of Iscove’s Modified Dulbecco’s Medium. In addition, the number of cells seeded per embryoid body was optimised to generate more uniform aggregates, improving handling and reducing variability during downstream differentiation.

However, a key distinction between the two differentiation strategies evaluated is the inclusion of Activin A and BMP4 in the A-HIC protocol. Addition of Activin A at the onset of differentiation likely facilitates a more directed exit from pluripotency by promoting SMAD2/3 signalling and activating transcriptional programmes associated with definitive endoderm commitment (McLean et al., 2007; Zhong et al., 2017, Cernilogar et al., 2019). Concurrently, BMP4 exposure mimics developmental cues involved in ventral foregut patterning and the acquisition of hepatic competence, directing endodermal progenitors toward a hepatic lineage rather than alternative fates such as pancreatic or intestinal lineages (Gouon-Evans et al., 2006; Shiraki et al., 2008; Cao et al., 2010). Together, these early morphogenic signals may support a more efficient transition through hepatoblast-like intermediates during the initial stages of differentiation. Therefore, another important insight gained from this study is that even though the marker expression trajectories did not show significantly differences, early cell fate-directed interventions in the A-HIC protocol promoted further hepatocyte maturation at endpoint.

Notwithstanding the variability observed across experiments, both differentiation protocols generated similar morphological and phenotypic characteristics. Cells differentiated with either protocol formed cobblestone-like clusters with well-defined borders, consistent with hepatocyte-like cells derived from pluripotent stem cells (Roelandt et al., 2010). Regions of dense clustering were interspersed with less populated areas, reflecting the EB-based differentiation strategy followed by flattening onto a culture surface, which was used to minimise necrotic core formation commonly observed in fully three-dimensional aggregates where nutrient and oxygen diffusion are limited (Gothard et al., 2010; McMurtrey, 2016). Although this approach improves nutrient accessibility, cultures did not form a uniform monolayer and instead stratified into multiple layers, but qualitative assessment by. immunofluorescence confirmed the presence of a range of cellular differentiation markers.

Consistent with transcript analyses, terminal cultures contained heterogenous populations expressing both hepatoblast (AFP) and mature hepatocyte markers (ALB), reflecting partial synchronisation of differentiation maturation. Nevertheless, both A-HIC and HIC protocols supported exit from pluripotency and progression toward a hepatic lineage. Hepatic marker expression stabilised by day 21, with no further maturation-associated changes detected at day 28. Prolonged culture primarily resulted in increased stratification rather than additional maturation, indicating that a 21-day differentiation period is sufficient to achieve the principal maturation features observed in this system. Therefore, this study has advanced a cost-effective reduction in method duration as well as of components without compromising on achieving more mature functional outcomes Additional functional assays further supported the hepatocyte-like phenotype of the differentiated cultures. Cells generated using both protocols could produce urea following ammonia stimulation, indicating the presence of active urea cycle metabolism. Similarly, both A-HIC and HIC cultures showed glutathione usage and sensitivity to inhibition of glutathione synthesis indicative of cellular redox capacity. Both protocols demonstrated the ability for uptake and excretory transport of indocyanine green (ICG) over time, consistent with functional hepatocyte-associated transport processes. Notably in several assays, differentiated cells performed more robustly than the HepG2 reference cell line. Despite variability across replicates, HepG2 cells failed to produce detectable urea, did not show glutathione synthesis inhibition under the tested conditions, and exhibited ICG uptake but limited ICG excretion. These observations indicate that, despite their widespread use, HepG2 cells do not fully recapitulate key hepatocyte metabolic and transport functions, whereas the differentiated cultures display several features consistent with hepatocyte-like activity.

In conclusion, the A-HIC protocol effectively directed murine embryonic stem cells toward a hepatocyte-like phenotype, supported by morphological, transcriptional, and functional characterisation. Differentiated cultures produced more consistent outcomes across independent experiments and supported a greater proportion of cells exhibiting mature hepatocyte characteristics. Importantly, this approach provides a relatively simple, cost-effective, and time-efficient differentiation method that may be readily adapted to more complex pluripotent systems, including hepatocyte differentiation from induced pluripotent stem cells and human iPSC lines, for applications in disease modelling and toxicology studies.

## Acknowledgments

The authors gratefully acknowledge Aaroy Hay (IRR Flow Cytometry and Cell Sorting Facility) for assistance with sample processing; Angus Comerford for provision of wild-type C57BL/6 mouse liver sections; Lynne Ramage for provision of indocyanine green; and Dr Shaden Melhem for provision of HepG2 cells. We thanks Dr David Duneau for his helpful early data discussion.

