## Supplementary Materials for "Development and characterisation of an optimised *in vitro* differentiation protocol for deriving hepatocyte-like cells from mouse embryonic stem cells"

### Supplementary Information

**Supplementary Table 1. *Pre-screening list of candidate mESCs hepatic differentiation protocols.*** The table summarises the hepatic differentiation protocols evaluated during preliminary optimisation, including published methods and study-specific modifications. For each protocol, the culture format and the key signalling factors with timing are reported and three independent biological replicates were performed priori to exclusion. Spontaneous differentiation was treated as a condition in its own right to assess whether hepatocyte-like cells could arise in the absence of exogenous signalling cues. Accordingly, three spontaneous conditions were included: EB-based differentiated in Iscove'medium (OIS), in DMEM (OC) and direct 2D differentiation in DMEM only (SD).

| Protocol source | Culture format | Key signalling factors and timing |
|---|---|---|
| Spontaneously Differentiated cells - OIS | EB-based+ plating in 2D | Spontaneously differentiated cells in Iscove's media |
| Hamazaki et al., 2001 | EB-based+ plating in 2D | aFGF (2 d) → HGF (6 d) → OSM/Dex/ITS (3 d) |
| Spontaneously Differentiated cells - SD | ES-E14TG2a "E14" plated in 2D | Spontaneously differentiated cells in DMEM media |
| Spontaneously Differentiated cells - OC | EB-based+ plating in 2D | Spontaneously differentiated cells in DMEM media |
| This study - A | EB-based+ plating in 2D | Chir99021 (1 d) → BMP4 (1 d) → BMP4 + aFGF (2 d) → HGF (6 d) → OSM/Dex/ITS (6 d) |
| This study - B | EB-based+ plating in 2D | Chir99021 (1 d) → BMP4 + aFGF (2 d) → HGF (6 d) → OSM/Dex/ITS (6 d) |
| This study - D | EB-based+ plating in 2D | Activin A + Chir99021 (1 d) → BMP4 (2d) + aFGF (4 d) → HGF (6 d) → OSM/Dex/ITS (6 d) |
| Zhou et al., 2009 | 2D using B6 & E14 as starting cell lines | Activin A (3 d) → SB/aFGF (5 d) → HGF/OSM/Dex (5 d) |
| Cao et al., 2010 | 2D using B6 & E14 as starting cell lines | SB/bFGF/BMP4 (6 d) → HGF (6 d) → OSM/Dex (3 d) |
| Rothová et al., 2016 | 2D using B6 & E14 as starting cell lines | Insulin/Activin A/EGF/FGF4 (6 d) *Anterior definitive endoderm/ventral foregut induction (not full hepatic differentiation; can be further differentiated into hepatocyte-like cells. |
| Pauwelyn et al., 2011 | 2D using B6 & E14 as starting cell lines | Activin A/Chiron (6 d) → BMP4/FGF2 (4 d) → FGF1/FGF4/FGF8b (4days) → HGF/Follistatin (6 d) |
| Pauwelyn et al., 2011 - with modifications - E14 | 2D using ES-E14TG2a "E14" as starting cell line | Activin A/Chiron (6 d) → BMP4/FGF2 (4 d) → FGF1/FGF4/FGF8b (4days) → HGF/Follistatin (6 d) |
| Pauwelyn et al., 2011 - with modifications - B6 | 2D using SCRC-1002 ES C57BL/6 "B6" as starting cell line | Activin A/Chiron (6 d) → BMP4/FGF2 (4 d) → FGF1/FGF4/FGF8b (4days) → HGF/Follistatin (6 d) |

### Supp Figure 1. Standard curve optimisation of qRT-PCR primers

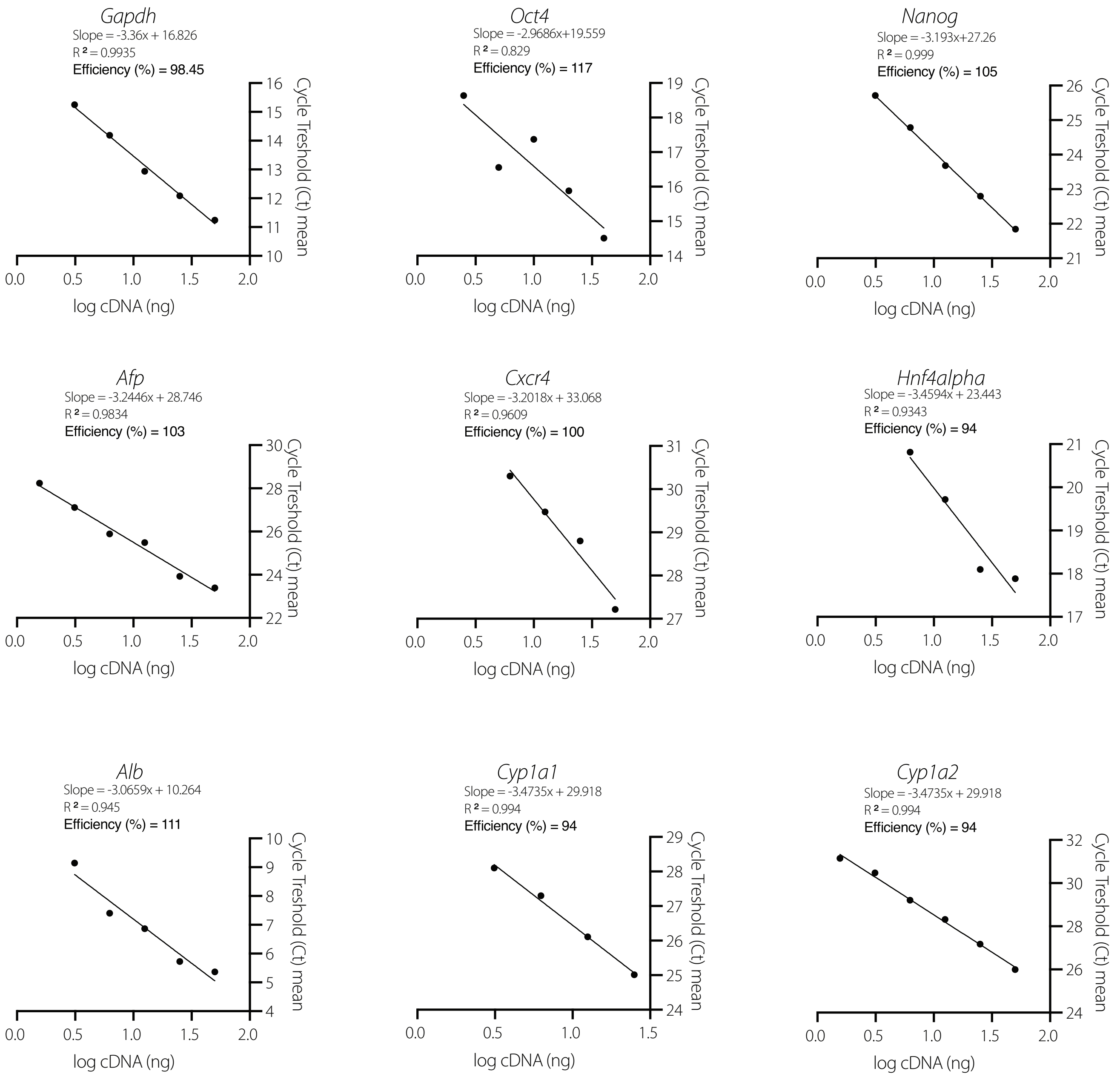

**Supplementary Figure S1:** ***Standard curve optimisation of qRT-PCR primers used for hepatic differentiation and metabolic characterisation.*** Standard curves were generated for primers targeting *Gapdh, Oct4, Nanog, Afp, Cxcr4, Hnf4α, Abl, Cyp1a1,* and *Cyp1a2.* cDNA was serially diluted two-fold across the indicated input range, with neat cDNA assigned an arbitrary value of 1.0 (log value 0). For *Oct4, Nanog,* and *Gapdh,* cDNA was derived from mouse embryonic stem cells (mESCs). For *Afp and Cxcr4*, cDNA was prepared from foetal mouse (C57BL/6) liver tissue collected at postnatal day 3. For *Hnf4a* and *Alb,* cDNA was generated from adult mouse (C57BL/6) liver tissue (28 weeks old). For *Cyp1a1* and *Cyp1a2,* cDNA was obtained from hepatocyte-like cells (HLCs) differentiated using the A-HIC protocol and exposed to 1 µM 3-methylcholanthrene (3MC) for 24 h to induce CYP expression. Quantitative RT-PCR was performed using each primer pair, and mean cycle threshold (Ct) values were plotted against log10 cDNA input. Linear regression was applied to derive standard curve slopes, which were used to calculate amplification efficiency according to the equation: Efficiency = ($10^{(-1/\text{slope}) - 1)}$ x 100). Correlation coefficients ($R^2$) are shown for each standard curve and indicate linearity across the dilution range

Supp Figure 2. Funnel-based selection of mESCs differentiation protocols for hepatic lineage for specification

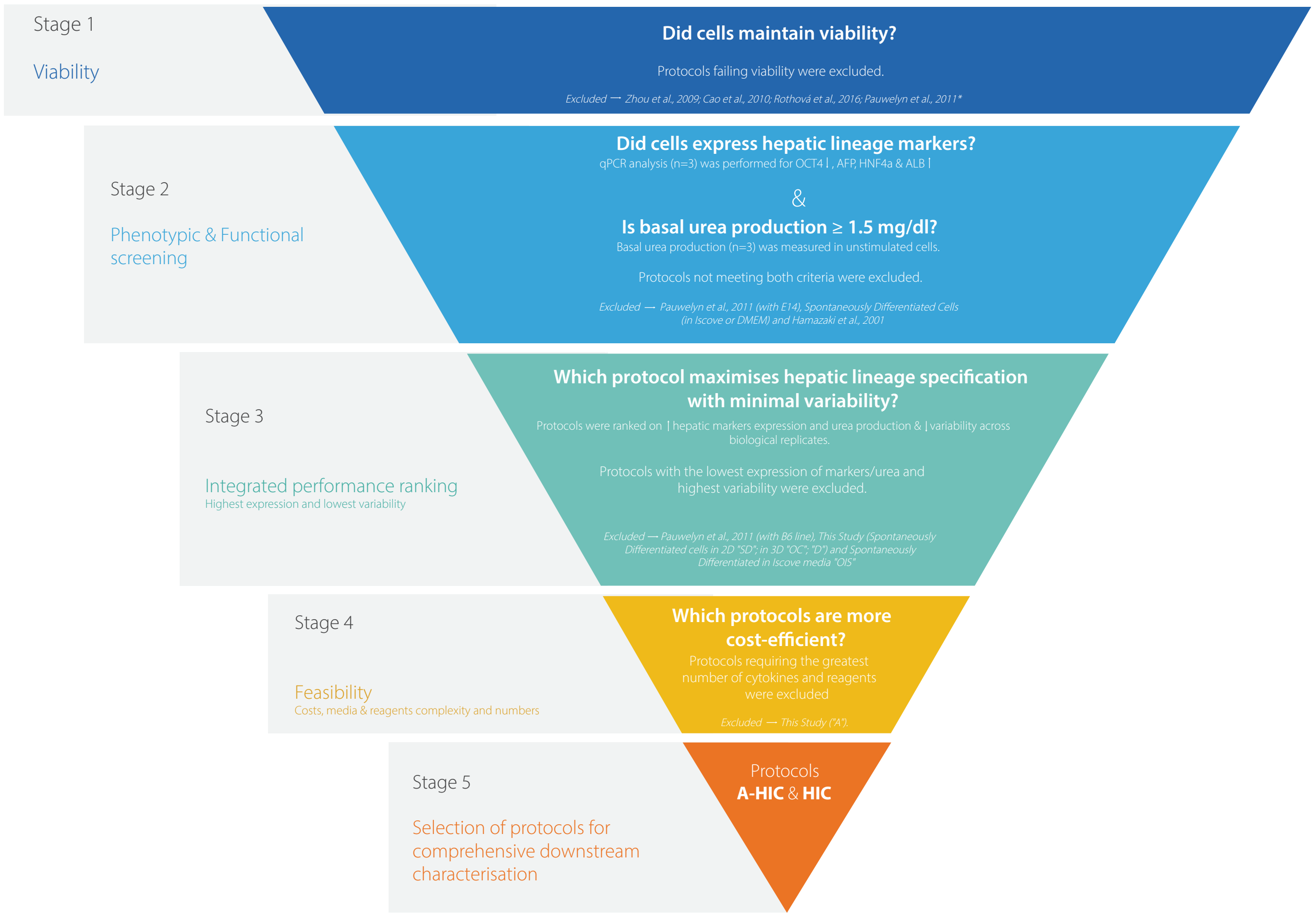

**Supplementary Figure S2. *Funnel-based selection of mESCs differentiation protocols for hepatic lineage specification.*** A multi-stage screening analysis was implemented to identify optimal protocols for hepatocytes-like cells differentiations. A total of 15 approaches (of which two being A-HIC and HIC) were first assessed. Stage 1 assessed cell viability, excluding protocols that failed to maintain viable cultures. Stage 2 evaluated hepatic lineage specification and functional competence based on qPCR expression of key markers (*Oct4, Afp, Hnf4alpha, Alb*) and basal urea productions, with protocols failing to meet both criteria excluded. Stage 3 involved an integrated performance ranking based on marker expression, urea production and variability across biological replicates, with lower performing protocols removed. Stage 4 assessed feasibility, excluding more complex protocols requiring additional cytokines/reagents where these did not confer measurable improvement over simpler conditions. This simplified, stepwise approach resulted in the selection of A-HIC and HIC protocols for comprehensive downstream characterisation (Stage 5).

Supp Figure 3. Immunofluorescence characterisation of hepatic markers.

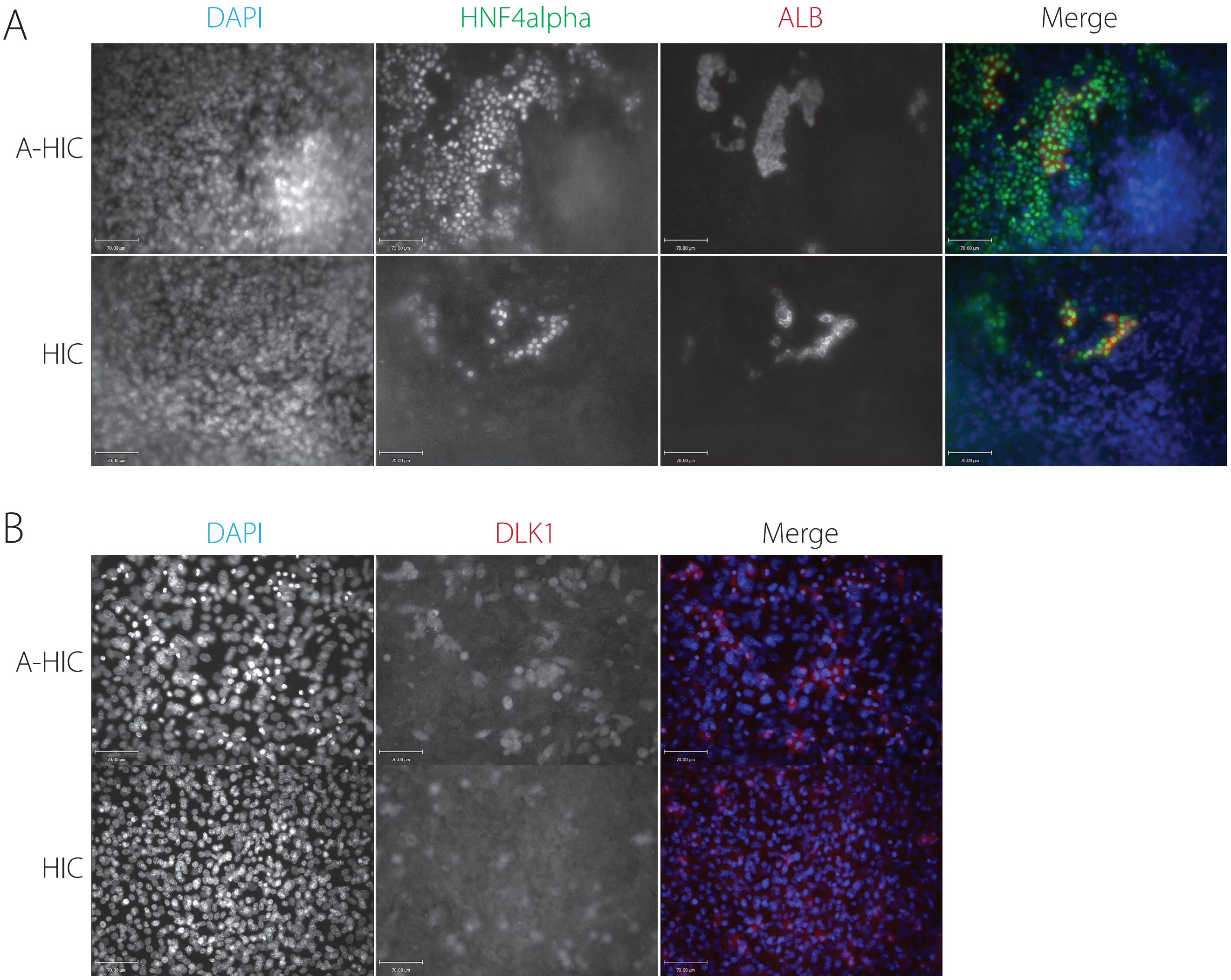

**Supplementary Figure S3.** ***Immunofluorescence characterisation of hepatic markers in A-HIC and HIC-derived cells*** Representative images show expression of the hepato-specific markers for both protocols. **A)** Co-staining for ALB (red, cytoplasmic) and HNF4alpha (green, nuclear) at lower magnification showing overall qualitative expression patterns across A-HIC and HIC-derived HLCs. Nuclei are counterstained with DAPI (blue). **B)** DLK1 staining in hepatocyte-like cells (HLCs) generated using A-HIC and HIC protocols showing a heterogeneous distribution with punctate, dot-like and cell surface-associated localisation. DLK1 is shown in red and nuclei are counterstained with DAPI (blue). Scale bars: 70 μm.

Supp Figure 4. qPCR analysis of metabolism specific CYPs over 28 days.

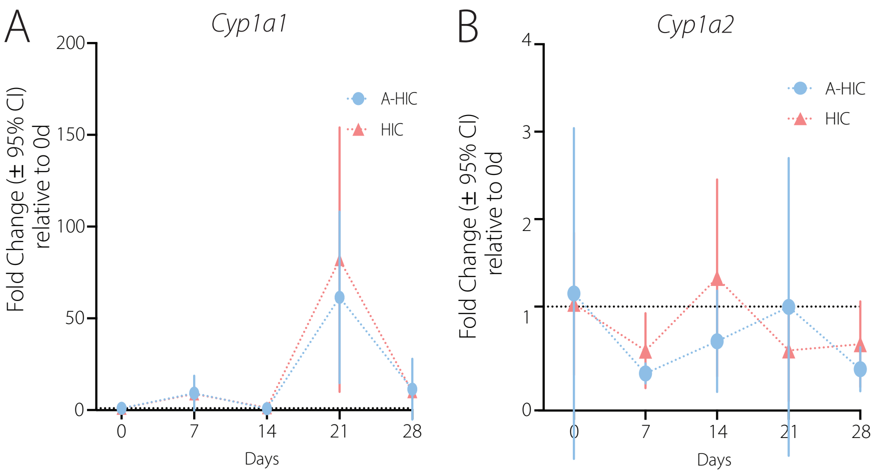

**Supplementary Figure S4.** ***qPCR analysis of metabolism specific CYPs over 28 days.*** **A-B)** Gene expression was measured at days 0, 7, 14, 21, and 28 in hepatocyte-like cells generated using both differentiation protocols **A)** Relative expression of baseline *Cyp1a1* main detoxifying gene **B)** Relative expression of un-baseline *Cyp1a2* main detoxifying. Expression of hepatic lineage markers *Hnf4α* and *Alb*. Data is presented as fold change relative to day 0 (mean ± 95% CI, *n* = 3 biological replicates). The red dashed line indicates baseline expression (fold change = 1).

### Supp Figure 5. Functional maturation and late-stage stability of HLCs during differentiation

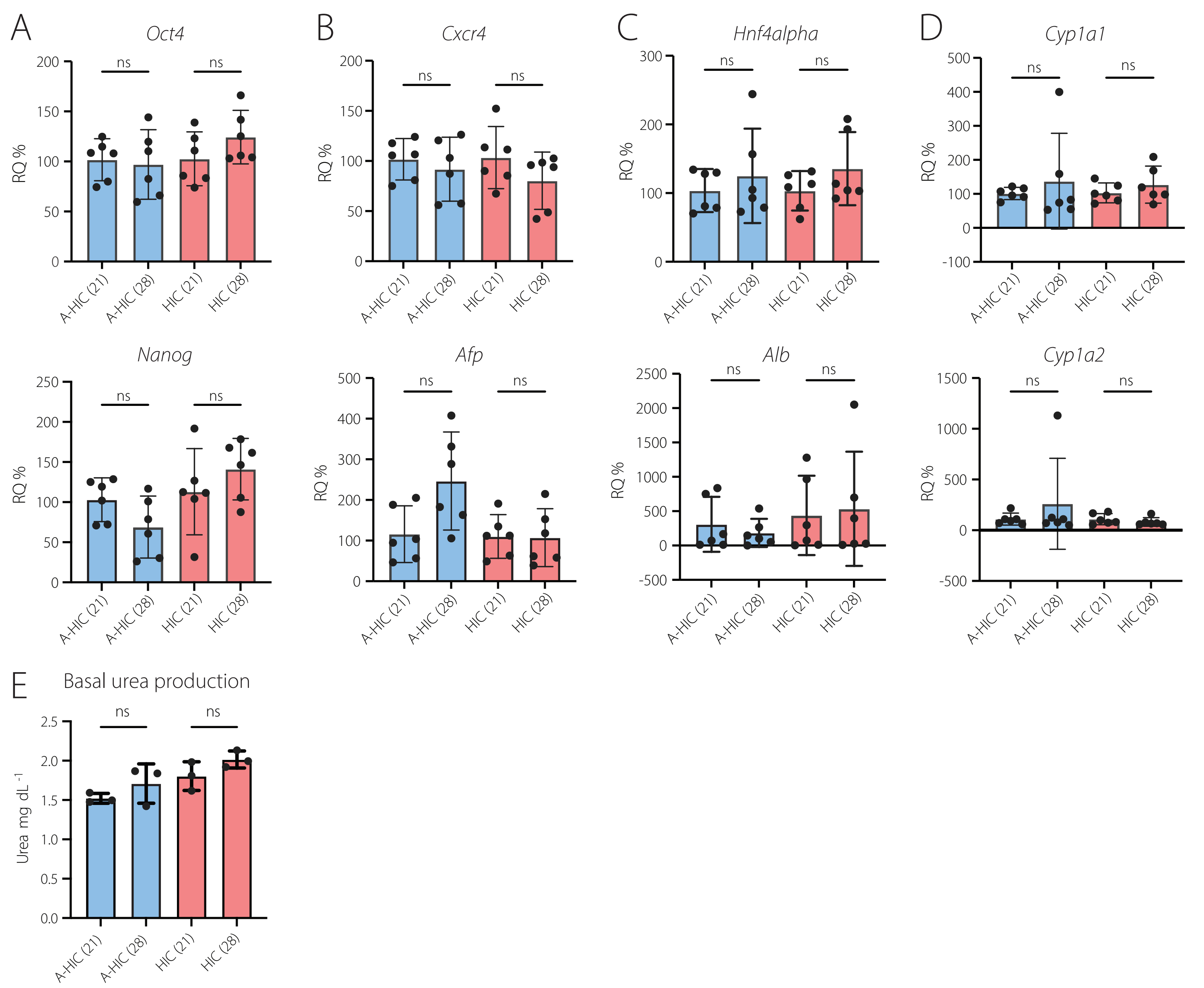

**Supplementary Figure S5.** ***Functional Maturation and Late-Stage Stability of Hepatocyte-Like Cells During Differentiation*** **(A-D)** Relative mRNA expression of selected panel genes in hepatocyte-like cells generated using protocols A-HIC and HIC at day 21 and day 28. **A**) Relative quantification of early pluripotency markers *Oct4*, *Nanog*. **B)** Relative quantification of intermediate meso-endodermal and hepatic progenitor markers *Cxcr4*, *Afp*. **C)** Relative quantification late hepatic lineage markers *Hnf4alpha*, *Alb***. D)** Relative quantification of detoxifying genes *Cyp1a1, Cyp1a2*. **E)** Basal urea production of HLCs generated with A-HIC and HIC at day 21 and day 28 measured in the absence of additional stimulation. For A-C values are shown as six individual biological replicates with mean ± 95% confidence intervals. Gene expression at day 28 was normalised to day 21 for each protocol and is presented as relative quantification (RQ). For D, values are shown as three individual biological replicates with mean ± 95% confidence intervals, illustrating inter-experimental variability. Analysis was performed using One-way ANOVA with Šídák's multiple comparisons test and it is indicated as *ns*, not significant.

Supp Figure 6. Validation of BSO-mediated inhibition

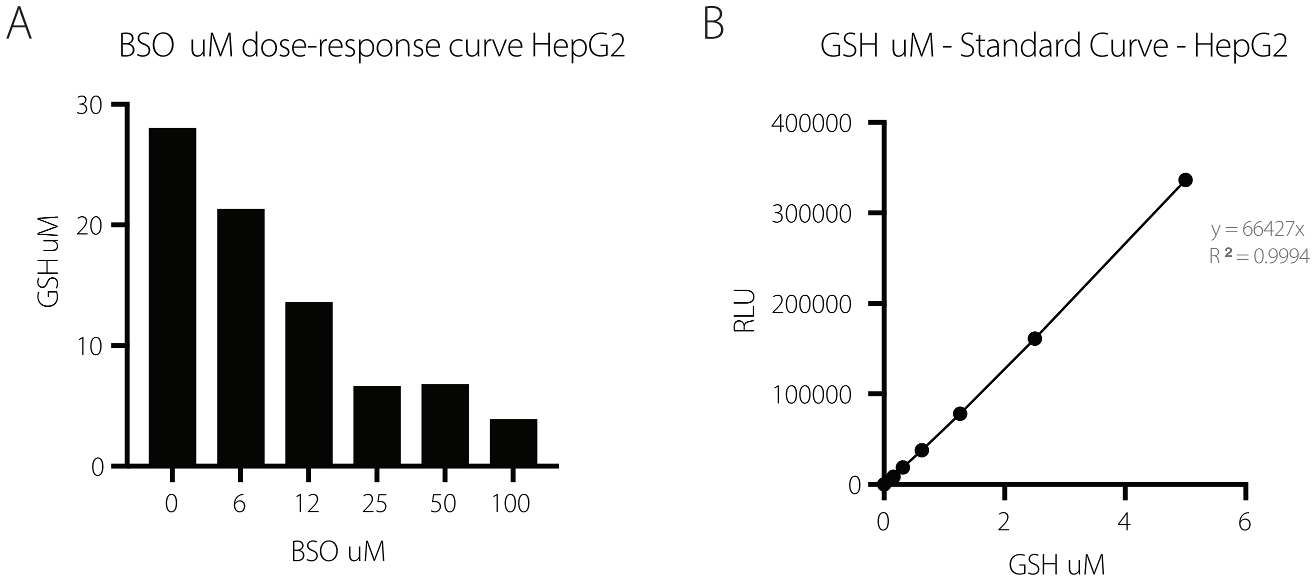

**Supplementary Figure S6.** ***Validation of BSO-mediated inhibition.*** **A**) Dose-dependent reduction in intracellular glutathione (GSH) levels depletion measured following exposure to increasing concentrations of buthionine sulfoximine (BSO); **B**) Representative standard curve generated for glutathione (GSH) quantification using HepG2 cell lysates, demonstrating a linear relationship between luminescence signal (RLU) and GSH concentration. Data are representative of assay validation experiments used to support subsequent functional analyses.

Supp Figure 7. Standard curves and viability after exposure to Indocyanine Green.

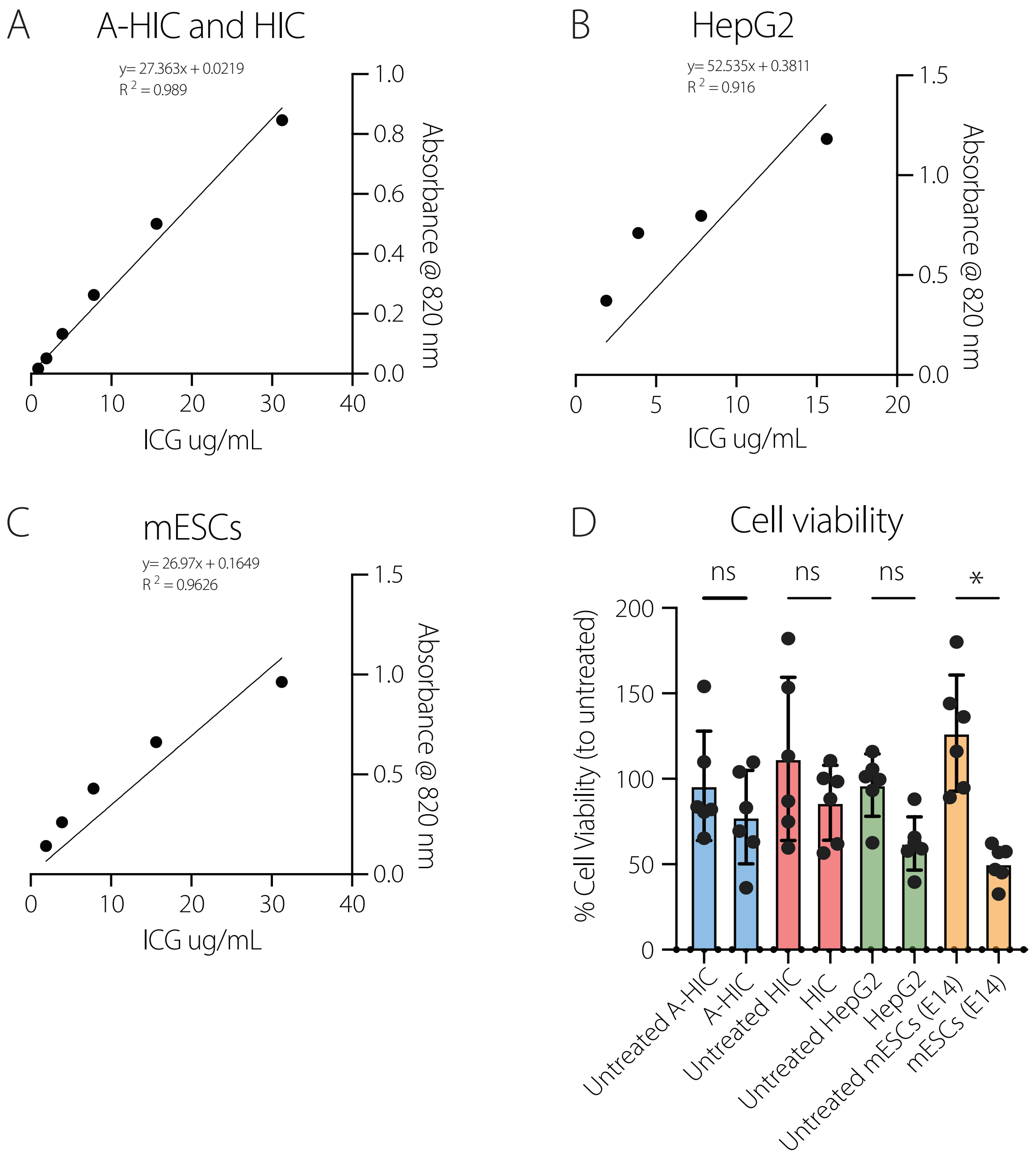

**Supplementary Figure S7.** ***Standard Curves and Viability after exposure to Indocyanine Green.*** **A)** Standard curve for indocyanine green (ICG) prepared in A-HIC and HIC media (DMEM with supplements and cytokines, excluding ITS due to insulin-related absorbance interference). **B)** Standard curve for ICG prepared in HepG2 culture media. **C)** Standard curve for ICG prepared in mouse embryonic stem cells culture media (including LIF). Serial dilutions (µg/mL) were used to derive linear regression models for absorbance-to-concentration conversion. **D)** Cell viability was assessed following ICG loading in hepatocyte-like cells (HLCs) generated using protocols A-HIC and HIC, alongside HepG2 cells and undifferentiated mouse embryonic stem cells (mESCs, E14). Viability is expressed as a percentage relative to untreated controls. Data are presented as individual biological replicates with mean ± standard deviation (SD). Analysis was performed using One-way ANOVA with Šídák's multiple comparisons test and it is indicated as *ns*, not significant or *p ≤ 0.05.

Supp Figure 8. IF analysis of hepatocyte marker retention for A-HIC HLCs on PET membranes

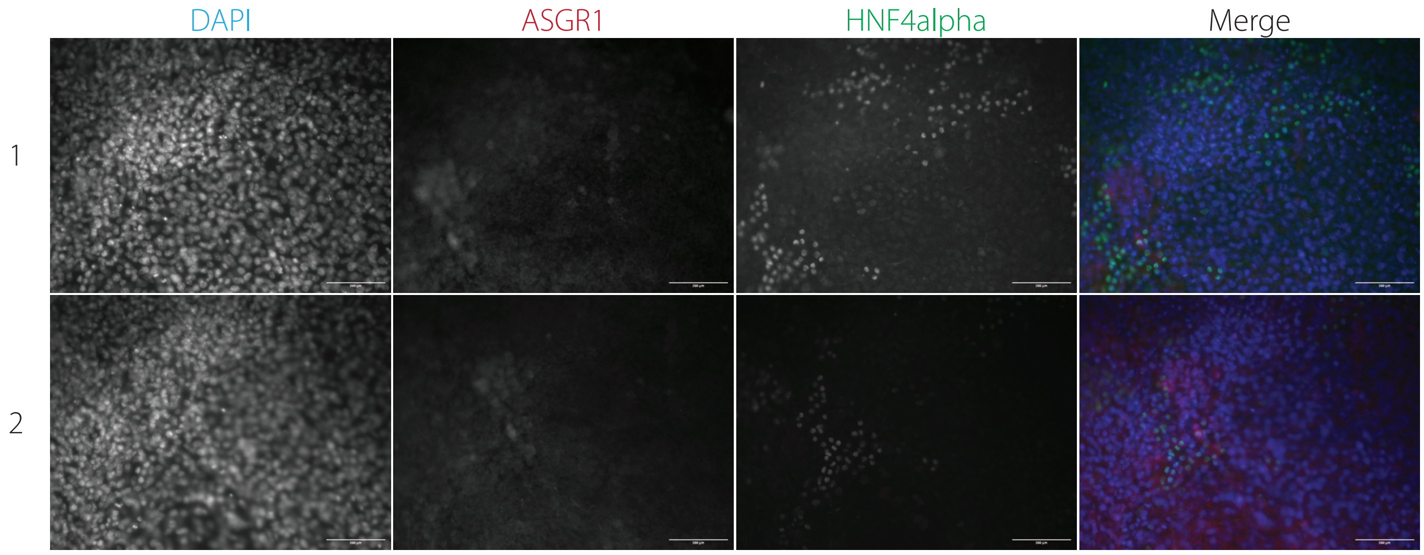

**Supplementary Figure S8.** ***Immunofluorescent analysis of hepatocyte marker retention for A-HIC HLCs on PET membranes.*** Representative immunofluorescence images of hepatocyte-like cells (HLCs) generated using protocol A-HIC following adhesion to transparent PET transwell membranes and stained with hepatic lineage markers HNF4a (green) and ASGR1 (red) with nuclear counterstaining by DAPI (blue). Scale bars: 70 µm. Images are representative of independent inserts.

Supp Figure 9. Flow cytometry gating strategy

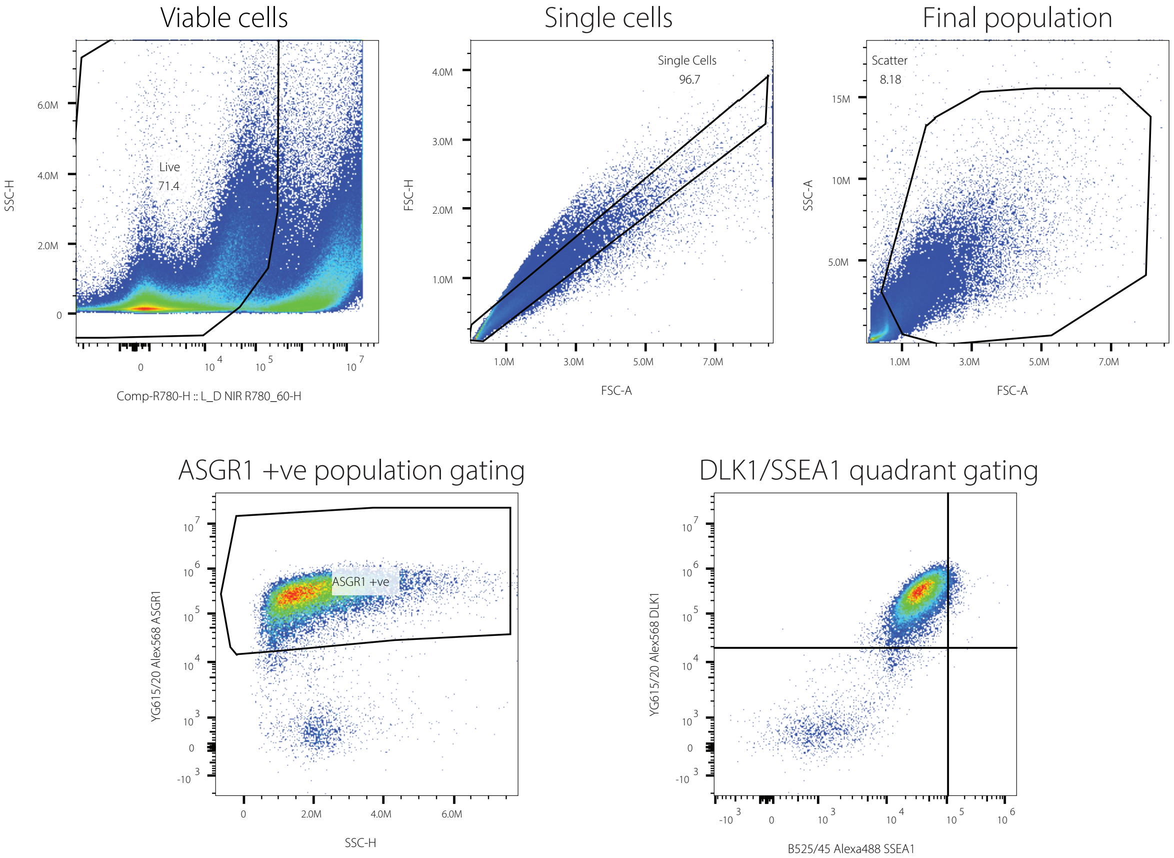

**Supplementary Figure S9.** ***Flow cytometry gating strategy and concatenated biological replicate analysis***. Representative gating strategy applied to all samples. Viable cells were first selected using Live/Dead Near-Infrared dye (x) versus SSC-H (y). Singlets were then gated using FSC-H versus FSC-A, followed by a scatter gate on SSC-A (y) versus FSC-A (x) to define the final population. For ASGR1 single staining, negative populations were defined using unstained, Live/Dead-only, ASGR1only, and secondary-only controls to establish the ASGR1$^{+}$gate. For DLK1/SSEA1 double staining, quadrants were applied with DLK1 on the y-axis and SSEA1 on the x-axis using the same controls.
